# Paper-and-pencil questionnaires analysis: a new automated technique to reduce analysis time and errors

**DOI:** 10.1101/2021.03.12.435109

**Authors:** Clovis Chabert, Aurélie Collado, Boris Cheval, Olivier Hue

## Abstract

**Background and Objective:** Questionnaires are essential tools in many scientific fields, including health and medicine. However, the analysis of paper-and-pencil questionnaires is time consuming, source of errors and expensive, limiting its use in large cohort studies. Computer-based questionnaires might be a valuable alternative but they may introduce bias, especially for sensitive questions, and they require programming skills. The aim of this study is to develop a reliable and adaptable open-source technique (i.e. LightQuest) to automatically analyse various types of scanned paper-and-pencil questionnaires with closed questions, including those with inverted scale.

**Methods:** To evaluate the usefulness of LightQuest, the time needed for 7 experimenters for manually code 10 sets of 4 frequently used questionnaires and the number of errors (i.e. reliability) were compared with the time and errors their made using LightQuest.

**Results:** LightQuest was twice as fast as the manual analysis, even though the time to create the reference model was taken into account (933s vs. 1935s, t(2)=8.81, p<0.001). Without model creation, the reduced analysis time was more pronounced, with an average of 2.77s.question^-1^ for the manual technique versus 0.55s.question^-1^ for LightQuest (t(2)=22.5, p<0.001). Moreover, during correction of the 5180 questions performed by the 7 experimenters, LightQuest made a total of 2 errors versus 46 with the manual technique (q(2)=4.53, p<0.05).

**Conclusion:** LightQuest demonstrated clear superiority both in terms of time and reliability. The script of this first open-source technique, which does not require programming skills, is downloadable in supplemental data and may become an asset for all studies using questionnaires.

## 1 Introduction

Questionnaires are ubiquitous in scientific fields ranging from psychology to epidemiology. They are used to assess numerous psychological health indicators in specific contexts, such as clinical care, as well as in more general contexts, such as epidemiological longitudinal follow-up. According to the Medline database, the keyword “questionnaire” appeared in 74,062 studies in 2016, corresponding to more than 5.8% of the total publications of that year. The occurrence of this keyword has increased over 10-fold in the past 50 years (0.51% of total publications in 1966), clearly indicating that the questionnaire has become an unavoidable tool in human research. However, their analysis is very time-consuming, especially in large cohort studies [1], and is a repetitive cognitively demanding task that is likely to generate errors despite the experimenter’s high degree of attention [2]. Previous studies, showed that almost all the spreadsheet studied showed errors despite great diversity in computerization methodology, and a visual corrections by experimenters [3, 4]. This introduce the concept of Garbage In/Garbage Out (GIGO), which express that the errors performed during the computerization of the data in spreadsheet software (i.e. garbage in) may lead to incorrect statistical analysis results (i.e. garbage out).

To decrease the errors from electronic transcription [4] and increase processing efficiency and reliability, computer-based versions of questionnaires have been developed [5–7]. Over the past 30 years, the development of communication tools and the computerization of results analysis have enabled large multi-centre studies that include thousands of subjects. However, computer-based questionnaires generally require strong programming skills; they may reduce the data quality due to the cognitive burden, the “yes” bias, the population recruitment [8, 9], and the time to computerization must be considered [1]. They may also introduce bias in specific populations such as adolescent cohorts, due to alteration of social inhibition [10], or elderly cohorts, who may be unfamiliar with computer use and therefore apprehensive [11]. Thus, computer-based questionnaires seem to distort results through disinhibition and a modification in social desirability [12, 13]. Furthermore, the differences in the results obtained with paper-and-pencil and computer-based questionnaires appear to be more pronounced for investigations seeking sensitive information (e.g. drugs, risky behaviours [14]), although several studies have shown that in some populations and under some circumstances, these two types of questionnaire do not produce different results [15, 16]. It should also be noted that computer-based questionnaires present disadvantages compared with paper-and-pencil questionnaires, notably in field experiments that require more organization and means. For example, for outdoor experiments a sufficient number of computers and an adequate power supply must be available, and in austere environments (e.g. rain, dust, low/high temperatures) computer fragility becomes an issue. Not least, most questionnaires were initially validated in paper format.

Overall, the automated analysis of paper-and-pencil questionnaires seems to be an interesting alternative cumulating both the advantages of computer-based questionnaires (i.e. time efficiency and error reduction) and paper-and-pencil questionnaires (i.e. logistics, cost, ecological task). To our knowledge, a few automated systems exist, but they have been designed only to correct multiple choice question (MCQ) tests and are not very adaptable as they have not been provided in an open source format [17, 18]. Several companies sell systems for automated analysis of paper-and-pencil questionnaires, but they are expensive and mainly destined for MCQ analysis in an educational context (e.g. OMR software or Exatech QCM).

Thus, the aim of this work was to develop an adaptable open source software to automatically analyse digitalized paper-and-pencil questionnaires with closed questions. The reliability (i.e. the number of errors) and efficiency (i.e. analysis time) of the technique were compared with the manual technique by analysing 4 frequently used questionnaires.

## 2 Material and Methods

### 2.1 Experimenters

Seven experimenters (29±4 years old) with 5.9±2 years of university education were recruited to analyse the questionnaires. Each experimenter analysed all the questionnaires manually and using the software in randomized order.

### 2.2 Analysed questionnaires

To compare the manual analysis with the software analysis, we used 10 sets of 4 well-known questionnaires, corresponding to a total of 740 questions. The questionnaires were: the Positive and Negative Affect Schedule (PANAS; [19]), the State-Trait Anxiety Inventory (STAI; [20]), the Profile of Mood States (POMS; [21, 22]), and the Rosenberg Self-Esteem Scale (RSES; [23, 24]). These questionnaires were chosen because they have been frequently used since their validation (respectively cited 27,562 times, 7,164 times, 9,659 times and 1,364 times) and, despite their lifespan, they are still used in many recent studies [25–28]. Although the RSES has been used less often than the others, its reversed scoring of some items makes it interesting for automated analysis because the reversed valence may increase the cognitive processes needed for analysis, which probably leads to increased analysis time and number of errors. The PANAS and the STAI are 20-item questionnaires with respectively 5 (ranging from 1 to 5) and 4 (ranging from 1 to 4) possible answers. The POMS consists of 24 questions on a 5-point scale ranging from 0 to 4. The RSES is a 10-item questionnaire with 4 possible answers (ranging from 1 to 4) and reversed valence for questions 2, 5, 6, 8 and 9.

### 2.3 Manual analysis

The 10 sets of each questionnaire were analysed in one session with a pause between the questionnaire types. The analysis time (in seconds) for each questionnaire corresponded to the time from the first question of the first questionnaire to the completion of answer digitalization of the 10^th^ questionnaire in a pre-established Excel matrix.

### 2.4 Automated analysis

The first step before starting the LightQuest script is to digitize an empty questionnaire, which will be used to create the model file, and the questionnaires to be analysed. For this study, digitization was performed with the charger of an ineo+554e printer (DEVELOP, Langenhagen, Germany), enabling us to copy all questionnaires in a single session. With this tool, the digitization time is approximately 1 second per questionnaire.

The automated analysis (i.e. LightQuest) has two main steps. First, the user has to create a model of the questionnaire to be processed (Figure 1A). In this step, which uses LightQuest_Model.m software for the analysis of a blank questionnaire, a model file is created and can be used every time the experimenter re-uses the questionnaire. Once this script is launched, a dialog box appears for the configuration of the questionnaire variables: number of items, response scale, presence of inverted items, and number of targets. Targets are the black rectangles visible on the questionnaire (see example of questionnaires used in this study in supplemental data “LightQuest.zip”) used for correcting the displacement (first target) and rotation (first and second targets) of the questionnaire, which may occur during digitization. Targets must be at the same height and have the same size (for this study we used 0.5cm x 0.5cm) and should be drawn on the questionnaire before printing. Once the user clicks on “OK”, the software asks the user to determine the approximate areas of the first and second targets, as described in “instructions for users” file (see supplemental data “LightQuest.zip”). The user then has to select all the questionnaire answers areas in one time to zoom and facilitate the selection of the question-by-question answer area. To obtain the best results, the selection rectangle must be focused on the centre of the answer area, with blank space all around. To help the user, the number of answers currently being parameterized is displayed at the top of the window. After selection of the answer area for the last question, a dialog box appears for the correction of the areas that the user estimates as wrongly selected. To finish creating the model, the user then simply clicks on “OK”.

**Figure 1:**
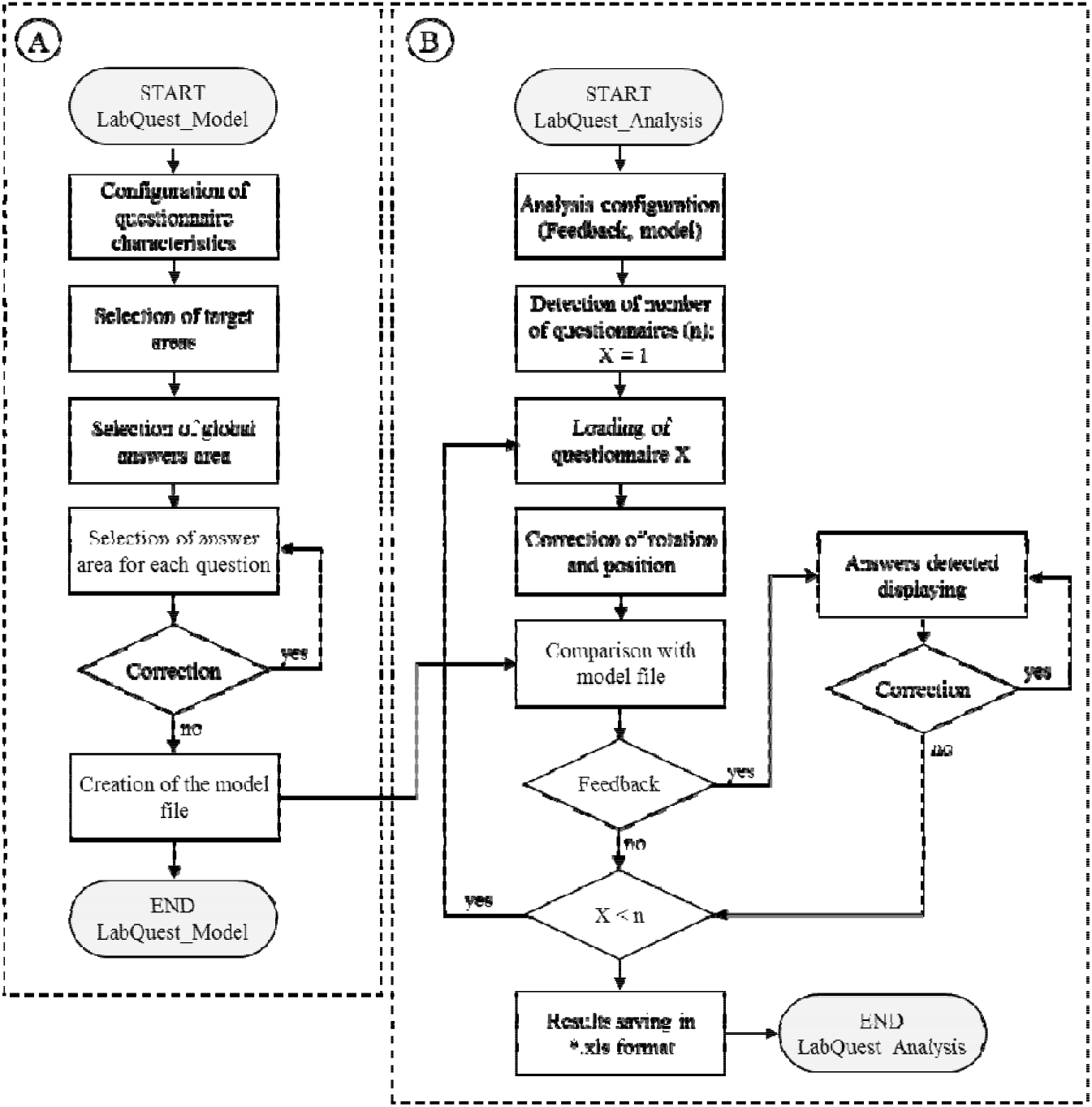
Main steps of LightQuest processing for questionnaire model creation (A) and analysis by experimenters of the completed questionnaires (B).

Second, the analysis of the questionnaires is performed by the LightQuest_Analysis.m file (Figure 1B). Once the script is run, a dialog box appears so that the appropriate model file can be selected to analyse the questionnaires, as well as the feedback level wanted during the analysis. If the feedback selected is “Yes”, each questionnaire is displayed with the detected answers, and the user can quickly verify and correct the analysis with an interactive interface in the case of multiple answers or no answer to a question (i.e: AF). Otherwise, selecting “No” means that the Excel matrix with the detected answers is directly generated by the script without steps of user verification and correction (i.e. AnF). For each participant of the study, 3 columns were implemented in the Excel file and saved in the Results folder. The first column corresponds to the detected answers, the second to the value attributed to the answer during model creation, and the third to the number of answers detected for each question. If inverted items are present in the questionnaire, the script automatically corrects the value attributed to the answer, in accordance with the configuration established during model creation. For ease of use, an explanation file describing the step by step use of the software is available in supplemental data (LightQuest.zip).

### 2.5 Statistical analysis

The analysis time, detected answers, and number of errors were collected for each technique and questionnaire. Analysis time (in seconds) per questionnaire and per question were assessed with an omnibus 2-way ANOVA according to the questionnaire (PANAS, STAI, RSES, PANAS) and technique (manually, automatically with feedback: AF to directly validate and correct the detected answers, and automatically with no feedback: AnF). To perform the omnibus 2-way ANOVA, data were corrected with Box-Cox transformation [29] to fit with normality law (Shapiro-Wilk test) and homogeneity (Levene test). Holm-Sidak *post-hoc* tests were performed to determine the detected effects. Because results did not show significant differences between questionnaires (results not shown), statistical analysis of the total number of errors with each technique (Figure 3) was performed with a Kruskal-Wallis 1-way ANOVA on ranks (technique factor), followed by a Tukey *post-hoc* test.

## 3 Results

### 3.1 Comparison of analysis times

Figure 2A presents the mean time taken by the 7 experimenters to analyse 10 copies of the PANAS, POMS, RSES and POMS questionnaires with the 3 techniques. Statistical analysis showed a main effect of both technique (F(2, 72) = 125.26, *p* < 0.001, partial *η2* = 0.74) and questionnaire (F(3, 83) = 4.66, *p* = 0.005, partial *η2* = 0.041), with no significant interaction between these 2 factors (F(6, 83) = 4.9e^-4^, *p* = 0.812, partial *η2* = 8.7e^-3^).

**Figure 2:**
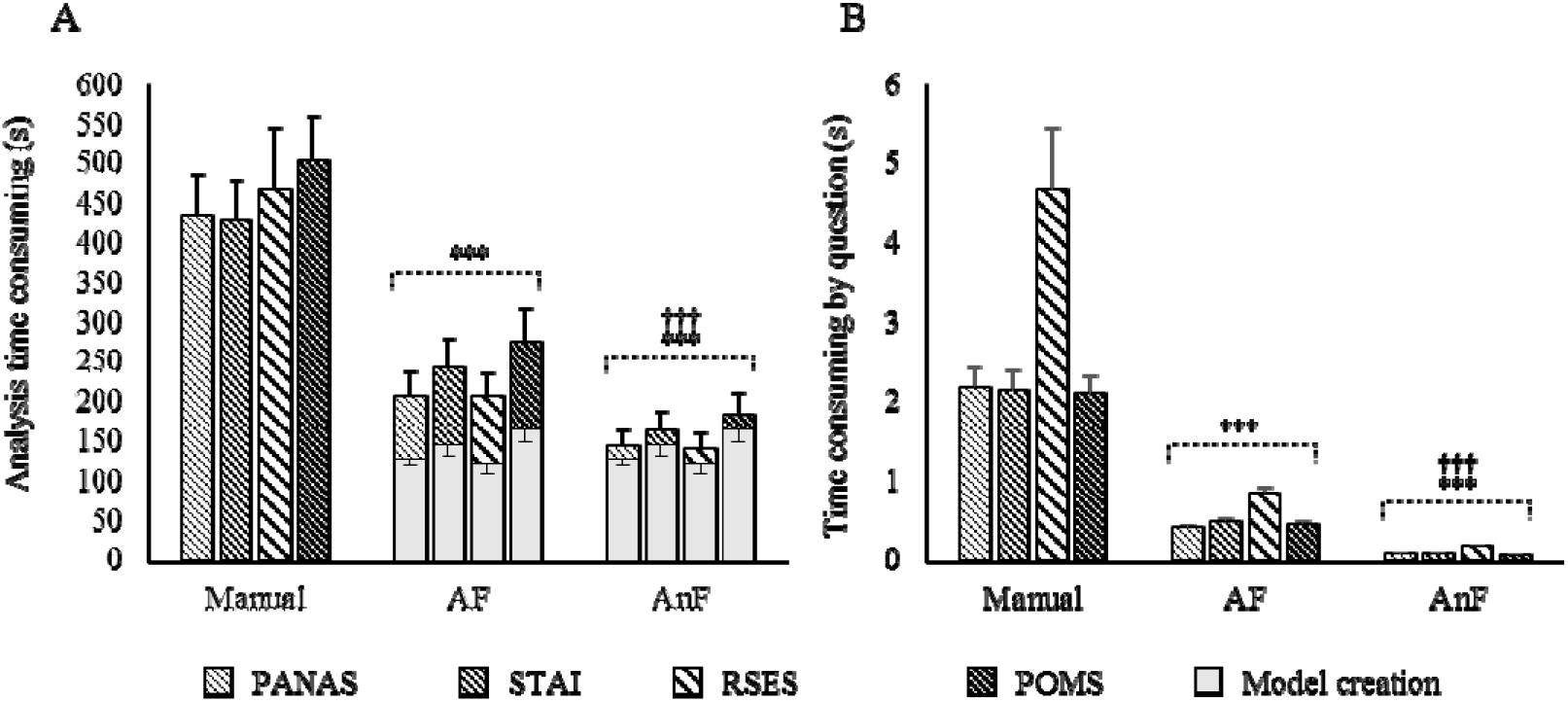
*Time needed for 10 questionnaires analysis (A) or for one question (B), according to the technique used. Mean±SEM; n = 7. AF: automatic + feedback; AnF: automatic with no feedback;* PANAS: *Positive and Negative Affect Schedule; STAI: State-Trait Anxiety Inventory; POMS: Profile of Mood States; RSES: Rosenberg Self-Esteem Scale. ***: diff. from manual analysis (p<0.001); †††: diff. from AF (p<0.001)*.

Comparison of manual *versus* AF and AnF showed a significant decrease in analysis time between 49% and 66% (manual *vs*. AF: t(2) = 8.81, *p* < 0.001; manual *vs*. AnF: t(2) = 15.79,*p* < 0.001). Comparison of AF and AnF revealed a significant 32% decrease in analysis time by technique: the time in AF condition was significantly longer than the time in AnF condition (t(2) = 6.98, p < 0.001). *Post-hoc* analysis of the questionnaire factor showed a significant difference between POMS (24 questions) *versus* the RSES (10 questions) and PANAS (20 questions) questionnaires (respectively t(3) = 3.35, *p* = 0.008; and t(3) = 3.06, *p* = 0.015).

Analysis time in relation to the number of questions in each questionnaire (Figure 2B) also showed a main effect of technique (F(2, 72) = 1083.42, *p* < 0.001, partial *η2* = 0.92) and questionnaire (F(3, 83) = 41.69, *p* < 0.001, partial *η2* = 0.053), with no significant interaction between these 2 factors (F(6, 83) = 0.57, *p* = 0.756, partial *η2* = 1.44e^-3^). Thus, the manual technique was 79% and 96% significantly slower than AF and AnF (respectively t(2) = 22.50, *p* < 0.001 and t(2) = 46.54, *p* < 0.001). AF and AnF were statistically different (t(2) = 24.03, *p* < 0.001), with the analysis time reduced over 80% with AnF compared with AF. Analysis of the questionnaire factor revealed that processing RSES was significantly longer than processing PANAS, POMS and STAI (respectively t(3) = 9.42, *p* < 0.001; t(3) = 9.39, *p* < 0.001; and t(3) = 8.46,*p* < 0.001).

### 3.2 Reliability of the analysis techniques

To test the reliability of the analysis techniques, we compared the total number of errors made by each experimenter when they corrected the questionnaires manually and with AF and AnF (Figure 3).

**Figure 3:**
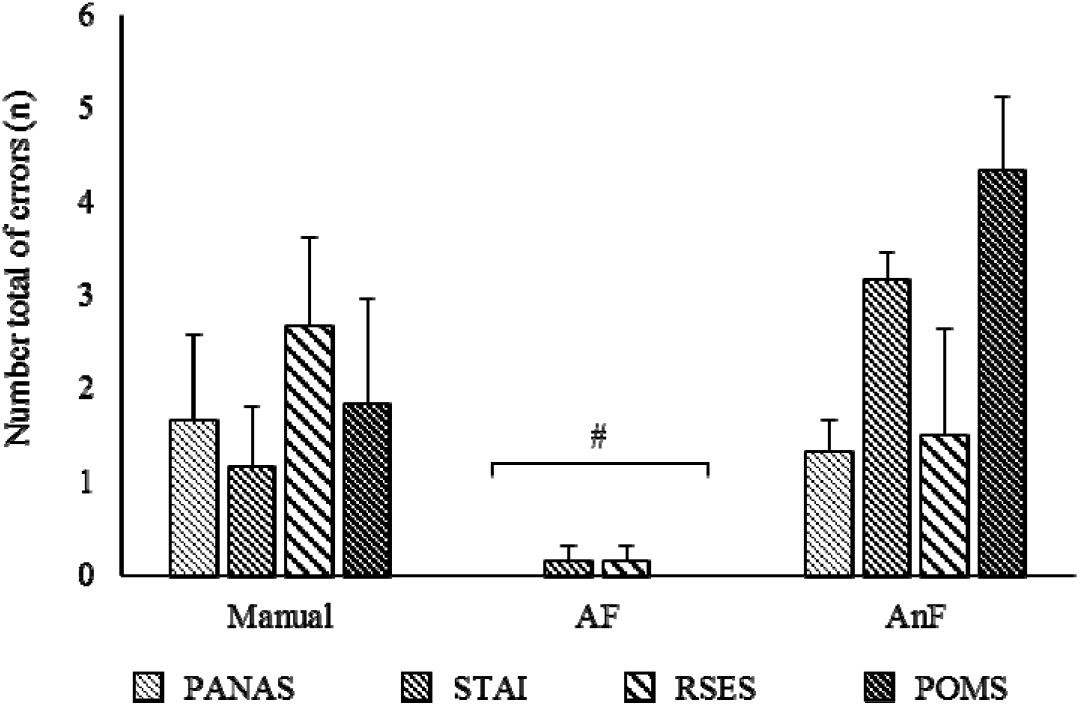
*Number of errors made by experimenters during analysis with the techniques. Mean±SEM; n = 7. AF: automatic + feedback; AnF: automatic with no feedback;* PANAS: *Positive and Negative Affect Schedule; STAI: State-Trait Anxiety Inventory; POMS: Profile of Mood States; RSES: Rosenberg Self-Esteem Scale. #: diff. from manual and AnF technique (p<0.05).*

A non-parametric 1-way ANOVA performed on the questionnaire factor found a significant difference (H(2) = 30.84, *p* < 0.001). Tukey *post-hoc* analysis showed that the number of errors with AF was significantly lower than with the manual (2 *vs*. 46 errors, q(2) = 4.528, *p* < 0.05) and AnF (2 *vs*. 67 errors, q(2) = 7.16, *p* < 0.05) techniques. No significant difference was observed between the AnF and manual techniques (67 *vs.* 46 errors, q(2) = 2.63).

## 4 Discussion

### 4.1 Main findings

Questionnaires are unavoidable tools in many research fields to better characterize study populations or evaluate psychological parameters, quality of life, fatigue, and so on. However, analysing a large number of questionnaires is expensive (if subcontracted) or very time-consuming, and mistakes are not uncommon. This study presents an efficient and reliable automated technique to analyse questionnaires with Matlab scripts (downloadable in supplemental data). Our results showed that the technique significantly decreased the number of errors (AF) and the time needed (AF and AnF) to process 4 widely used questionnaires, suggesting that it is a potential asset in all studies using paper-and-pencil questionnaires. Furthermore, this technique enables users to configure new questionnaires, thereby making it highly adaptable.

### 4.2 Comparison with manual technique

The automated technique with user feedback (AF) reduced the time needed to manually analyse 10 sets of 4 questionnaires (1835s *vs.* 933s) by half and appeared to be time-efficient from the 5^th^ questionnaire. At the 20^th^ questionnaire, the AF technique was 9 to 11min faster and saved effective-time analysis work. Furthermore, this benefit included the time needed to create a model, corresponding to approximately 60% of the processing time, whereas the model can be re-used as long as the questionnaire is not changed. Thus, once the model is created, the technique is at least 4.4 to 5.5 times faster than manual analysis, and this is without taking into account the decreased efficiency in analysis due to the experimenter’s fatigue during a repetitive task [30].

In addition to the time saved, the automated technique with feedback sharply decreased the number of errors. Indeed, the number of errors occurring during analysis was divided by 23 for the AF technique compared with the manual analysis. For the RSES, the number of errors with manual analysis was identical to those for the other questionnaires, despite differences in the number of questions (only 10 for RSES *vs*. 20-24 for the others). Thus, the number of errors per question was doubled for the RSES, probably due to the reverse-coded questions that increased the task complexity. With this technique, inverted items are configured during model creation and are automatically taken into account by the script during analysis.

### 4.3 Interest of the automatic with no feedback (AnF) option

An option was also developed to directly generate the results table without user feedback and correction (AnF). With this option, computer processing took less than 2s per questionnaire, whereas the AF technique needed 11s. However, using this option resulted in a gain of only 1-2min in analysis time for each set of 10 questionnaires and sharply increased the total number of errors compared with AF (2 vs. 67 errors for 280 questionnaires). Furthermore, although the difference appeared to be non-significant in this study, the number of errors was multiplied by almost 1.5 compared with manual analysis (67 errors *vs.* 46 for 280 questionnaires). Thus, we recommend using this option only in very specific situations after careful verification of the model quality. A post-analysis correction is also possible directly in the Excel table generated by the script, facilitated by the annotation of the number of answers detected for each question in the third column. Nonetheless, this correction mode is not advisable because it appears to be more time-consuming than the user interface we developed to directly validate the software analysis (data not shown). Interestingly, the low number of errors obtained with the model created by beginners shows the ease of handling our technique (experimenters have only one created model to become used to).

### 4.4 Strengths and weaknesses

To our knowledge, LightQuest is the first open source script available for the analysis of paper-and-pencil research questionnaires, and the possibility of creating new model files makes it an usable tool for all existing and future questionnaires. Furthermore, LightQuest’s graphic interface is user-friendly, decreasing the cognitive load and thereby decreasing the questionnaire analysis time and the number of errors. LightQuest appears to be particularly efficient for analysing questionnaires with an inverted scale, probably because it requires fewer cognitive resources compared with manual analysis. Last, the 2 black targets help to correct the rotation and displacement of the questionnaire during computerization, which makes LightQuest functional with all printer chargers or commercial scanners.

However, to maintain sufficient picture quality, the questionnaires need to be printed from a computer and photocopiers should not be used. For already completed questionnaires, the absence of 2 black targets reduces LightQuest’s accuracy. Nonetheless, an option is available (i.e. Black target = 0) to analyse questionnaires without these targets, but the low accuracy implies more manual corrections by the LightQuest user. The AnF option decreases the analysis time but increases the number of errors. However, these errors are mainly because LightQuest is unable to identify mistakes made by study participants (e.g. 42 cross- outs out of the 160 questionnaires corrected during this study) and the software makes artefactual detections when participants exceed the response box. Thus, this option is only recommended for questionnaires with considerable distance between answer areas and after visual validation of the created model quality. Last, LightQuest is not encoded to analyse visual analogue scales, but this function could easily be implemented in its open source code by a user with programming skills.

## 5 Conclusion

The open source script proposed in this study considerably reduces the analysis time and the number of errors on paper-and-pencil questionnaires. Thus, using LightQuest for questionnaires based studies may reduce cost, allow inclusion of larger cohorts and decrease the errors of interpretation due to mistake during the manual electronic transcription of data. LightQuest is adaptable to any questionnaire with closed questions by adding black targets, and no programming skills are required. To our knowledge, this technique is the only one offering automated analysis of research questionnaires, which is why it could become an asset for large cohort studies in many fields of investigation.

## Supporting information

LightQuest_script_package

## Abbreviations

AF: Automatic with Feedback
AnF: Automatic with no Feedback
GIGO: Garbage In/Garbage Out
MCQ: multiple choice question
PANAS: Positive and Negative Affect Schedule
POMS: Profile of Mood States
RSES: Rosenberg Self-Esteem Scale
STAI: State-Trait Anxiety Inventory

## Acknowledgement

This project was supported by a « Programme Opérationnel - Fonds Européen de Développement Régional » (PO-FEDER) grant. B.C. is supported by an Ambizione grant (PZ00P1_180040) from the Swiss National Science Foundation (SNSF).

## Author Contribution Statement

C.C. and A.C. participated equally in the design, measurement, analysis and redaction of this work. B.C. and O.H. contributed to the data analysis and the redaction of this work.

## Conflict of interest

No conflict of interest is declared by the authors.

## Notes

### Competing Interest Statement

The authors have declared no competing interest.

