## Supplementary material for "Paper-and-pencil questionnaires analysis: a new automated technique to reduce analysis time and errors": LightQuest_script_package: Instructions for users.pdf

### First Use

In the Matlab folder (generally: "C:\Users\Name\Documents\Matlab\") extract the file « LightQuest.zip » downloadable in supplemental data. This file contains the folders named "Files", "Scripts", "Results" and "Models".

### Creation of the Model file of your questionnaire

If you are using a new questionnaire, you have to create the model file before the analysis.

1) After copying the blank questionnaire you used in the folder "Files", launch the script "LightQuest_Model.m" with the Matlab interface.
2) Complete all fields of the window which has just appeared and click on "ok".
3) Approximately select the first and second target of the questionnaire as describe in figure 1A of the article.

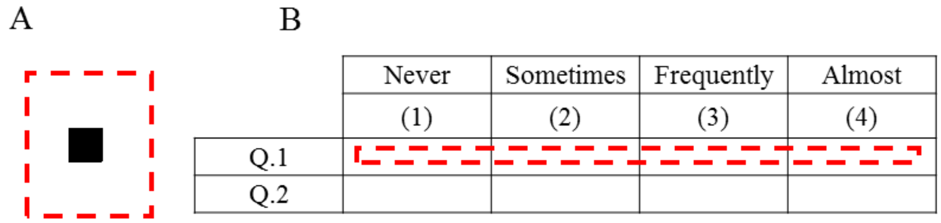


*Supplemental figure 1: Example of the selection of a target area (A) and an answer area (B). The red dashed rectangles are the selections made by the user.*

4) Select the global area containing all the answers of your questionnaire to zoom on it and facilitate the selection of each answers' areas one by one.
5) For each question, select the area were the subjects of your study are likely to answer. As describe in the figure 1B of the article, don't forget to center your selection box and to maintain margins around the area to not include the lines around the question.

Rq: For helping you in your task, the number of the question in progress is displayed at the top of the figure.

6) Once all answers areas were selected, the script allows you to correct those you estimate poorly selected. Just tip the number of the question in the window and click "ok".
7) When user is satisfied by all the selected areas, just click on "ok" (with "0" in the window field) to finish the script and save the model file. The file is saved in the folder "Models" and named as the empty questionnaire file you used.

   Rq: If you used several computer for the questionnaires analysis, you just need to copy this file in folder "Models" of the others computers.

**Analyze of your questionnaires**

1) First copy the files you want to analyze in the folder "Files" and suppress the blank questionnaire you have used to create the model.
2) Launch the script "LightQuest_Analysis.m" with the Matlab interface and complete all fields.
3) Analyzed questionnaire is displayed with bounding boxs on the answers detected by the script. Green box appears if one answer is detected, red box appears if 2 or more answers are detected and blue box appears if no answer is detected for the question.
4) If a visual validation of analysis by user was selected, for each misdetected answers the user may correct it by clicking on the area of the right answer. In case of no right answer exist, user have to click in the grey area bounding the figure to go next.
5) Once the misdetected answers by the script are corrected by user, it's also possible to manually change the answer for each question (by entering the number of the question whose answer have to be modified). When satisfied by all the areas selected, just click on "ok" (with "0" in the windows field) to go to the next questionnaire.

   Rq: The number of the questionnaire currently analyzed and the total number of questionnaires are displayed at the top of the figure.

6) After analysis of the last questionnaire, results are saved in the folder "Results" in an excel file named "*Model Name*_Analysis.xls". If a file with this name already exists, results are saved under the name: "*Model Name*_Analysis _Copy(n).xls".
