## Supplementary material for "Paper-and-pencil questionnaires analysis: a new automated technique to reduce analysis time and errors": LightQuest_script_package: LightQuest_Analysis.m

```
clear all
clc
tic


%%%%%%%%%%%%%%%%%%

prompt = {'Filename of the model (without ".mat"):','Visual validation of analysis by user (Yes or No):'};
dlg_title = 'Input';
num_lines = 1;
defaultans = {'Model_POMS','Yes'};
answer = inputdlg(prompt,dlg_title,num_lines,defaultans);


MCQ_Model = answer{1};
V_Feedback_Temp = answer{2};

if V_Feedback_Temp == 'Yes';
V_Feedback = 1;
else
    V_Feedback = 0;
end

%FOLDERS
Files_Folder = '../Files/'; % XML Folder
Model_Folder = '../Models/'; % Matrix Folder (with file created by 'a_Tracking_script.m')
Output_Folder = '../Results/'; % folder containing the XLS result files

%%%%%%%%%%%%%%%%%%%%%%%%%%%%%
disp('>> TESTING PARAMETERS');
if ~exist(Files_Folder,'dir');
    uiwait(errordlg('Files folder not exists. Please check it.'));
elseif ~exist(Model_Folder);
    uiwait(errordlg('Model folder not exists. Please check it.'));
elseif ~exist(Model_Folder);
    uiwait(errordlg('Results folder not exists. Please check it.'));    
end
List_Model = dir([Model_Folder MCQ_Model '.*']); 
List_Files = dir([Files_Folder '*.jpg']); 
List_Model_JPG = dir([Files_Folder MCQ_Model '.jpg']); 

if length(List_Model) < 1;
    uiwait(errordlg('Model file not found, please check if present or check the file name of the model and run again the script'));
elseif length(List_Files) < 1;
    uiwait(errordlg('Questionaries files not found, please check if present in Files folder and run again the script'));  
elseif length(List_Model_JPG) == 1;
    uiwait(errordlg('Picture used to create the model is still present in the Files folder, please suppress it and run again the script'));     
    error
elseif length(List_Files) < 1;
    uiwait(errordlg('Model file not found, please check if present or check the file name of the model and run again the script')); 
elseif ~((V_Feedback == 0) | (V_Feedback == 1));
    uiwait(errordlg('Feedback has to be "Yes" or "No". Please change Feedback entry and run again script.'));
    error
else
    disp('Parameters ok. Running script goes on.');
end


%%%%%%%%%%%%%%%%%%
load([Model_Folder '' MCQ_Model '.mat']);


%Acquisition of variables
Nb_Page = [Model(1,1,2)];
Nb_Question = [Model(2,1,2)];
Nb_Choice = [Model(3,1,2)];
Values(1,:)= Model(4,:,2);
Nb_Inv_Question(1,:) = Model(5,:,2);
Edge = (1.8*[Model(6,1,2)]);
Nb_Target = [Model(7,1,2)];
Auto_Thresh = 3;
Model2 = Model (:,:,:);

s=1;
Auto_Thresh_Save = zeros(Nb_Question,1);


% Detection of the number of questionnaries
Files_Detection = dir([Files_Folder '*.jpg']);
Nb_Files = length(Files_Detection(:,1));


for z = 1:Nb_Files;

File_Name = strcat(Files_Folder,Files_Detection(z).name);
Picture = imread([Files_Folder '' File_Name]);

%transformation of Picture in binary
Picture = im2bw(Picture,0.9);


%Correction of the rotation
if Nb_Target == 2;
    for Ar= 1:2;
        Pic = Picture(Model([6+Ar],4,2):Model([6+Ar],5,2),Model([6+Ar],6,2):Model([6+Ar],7,2));
        Pic = imcomplement(Pic);
        Pic = bwareaopen(Pic,300);
        bw = bwlabel(Pic, 4);
        stats1 = regionprops(bw, 'Centroid','Area');

        Rot_cor(Ar,1)= stats1(1).Area;
        Rot_cor(Ar,2:3)= stats1(1).Centroid;
        Rot_cor(Ar,2)=Rot_cor(Ar,2)+Model([6+Ar],6,2);
        Rot_cor(Ar,3)=Rot_cor(Ar,3)+Model([6+Ar],4,2);
    end


    %Rotation of the Picture
    Rot_cor(3,1) = abs(Rot_cor(1,2)-Rot_cor(2,2));
    Rot_cor(4,1) = abs(Rot_cor(1,3)-Rot_cor(2,3));
    S_Pic = size(Picture);
    Picture = imrotate(Picture,round([atand(Rot_cor(4,1)/Rot_cor(3,1))],3));
    Mrot = ~imrotate(true(S_Pic),[atand(Rot_cor(4,1)/Rot_cor(3,1))]);
    Picture(Mrot&~imclearborder(Mrot)) = 1;
end

if Nb_Target>0;
    %Detection of the new centroid of the first target
    Pic2 = Picture(Model(7,4,2):Model(7,5,2),Model(7,6,2):Model(7,7,2));
    Pic2 = imcomplement(Pic2);
    Pic2 = bwareaopen(Pic2,300);
    bw = bwlabel(Pic2, 4);
    stats2 = regionprops(bw, 'Centroid');
    Center = stats2(1).Centroid;
    Center(1,1)= Center(1,1)+ Model(7,6,2);
    Center(1,2)=Center(1,2)+ Model(7,4,2);

    %Comaparison with thus detected on the model
    Center(1,1) = round(Center(1,1) - Model(9,2,2));
    Center(1,2) = round(Center(1,2) - Model(9,3,2));

    %increments of the new coordinates of answer's areas
    Model(10,4:5,2) = Model(10,4:5,2) + Center(1,2);
    Model(10,6:7,2) = Model(10,6:7,2) + Center (1,1);
end

% Picture question area
Picture2 = Picture(Model(10,4,2):Model(10,5,2),Model(10,6,2):Model(10,7,2));

%matrix creation
Results = zeros (Nb_Question, 50);
Results0 = zeros (Nb_Question, 5);
Results2 = zeros (Nb_Question, 5);

%Analysis of each Question
for i = 1:Nb_Question;
Nb_Ans = 2;
count=1;
 while Nb_Ans ~= 1 && count <=30;
  for q =1:Nb_Choice
    Results0(i,q)=nnz(~(Picture2([Model(i,2,1)]:[Model(i,3,1)],[Model(i,[q+3],1)+Edge]:[Model(i,[q+4],1)-Edge])));
    Results(i,q) = Results0(i,q)-Model(i,[q+4+Nb_Choice],1);
  end
  
  Nb_Ans = sum(Results(i,1:Nb_Choice) >= Auto_Thresh);
  
  % Auto_Threshold adjustements
  if Nb_Ans > 1 && Auto_Thresh < 300;
  Auto_Thresh = Auto_Thresh+5;
  end
  
  if Nb_Ans == 0 && Auto_Thresh > 50;
  Auto_Thresh = Auto_Thresh-5;
  end
 count = count+1;
 end

 %Save of the Auto_Thresh Value
 Auto_Thresh_Save(i,1) = Auto_Thresh*0.975;
 
  max_num=max(Results(i,1:Nb_Choice));
 
  if max_num>Auto_Thresh*0.975;
    [row, col] = find(ismember(Results(i,1:Nb_Choice), max_num));
    Results2(i,2) = col(1,1);
  end
   Results2(i,4) = sum(Results(i,1:Nb_Choice) >= Auto_Thresh*0.975);
   


end

%%%%%Feedback%%%%%%%
if V_Feedback == 1;
    n2=1;
    n3=1;
    Col = zeros(Nb_Question*2,3);
Feed_Mat = zeros([Nb_Choice*Nb_Question+50],8);
for i2=1:Nb_Question;
[Xv Yv] = find (Results(i2,1:Nb_Choice) >= Auto_Thresh_Save(i2,1));
v = nnz(Xv);
%implement of answer(s) detected in Feed_Mat
if v==1;
    Feed_Mat(n2,2) = Model(i2,2);
    Feed_Mat(n2,1) = Model(i2,[Yv(1,1)+3]);
    Feed_Mat(n2,4) = Model(i2,3)-Model(i2,2);
    Feed_Mat(n2,3) = Model(i2,[Yv(1,1)+4])-Model(i2,[Yv(1,1)+3]);
    Feed_Mat(n2,5:7) = [0 1 0];
    n2=n2+1;
end
%implement of Feed_Mat if no answer was detected
if v == 0 ;
    Feed_Mat2(n3,2) = Model(i2,2);
    Feed_Mat2(n3,1) = Model(i2,4);
    Feed_Mat2(n3,4) = Model(i2,3)-Model(i2,2);
    Feed_Mat2(n3,3) = Model(i2,[Nb_Choice+4])-Model(i2,4);
    Feed_Mat2(n3,5:7) = [0 0 1];
    Feed_Mat2(n3,8) = i2;
    n3=n3+1;
end

if v >1;
for n=1:v;
Feed_Mat2(n3,2) = Model(i2,2);
Feed_Mat2(n3,1) = Model(i2,[Yv(1,n)+3]);
Feed_Mat2(n3,4) = Model(i2,3)-Model(i2,2);
Feed_Mat2(n3,3) = Model(i2,[Yv(1,n)+4])-Model(i2,[Yv(1,n)+3]);
Feed_Mat2(n3,5:7) = [1 0 0];
Feed_Mat2(n3,8) = i2;
n3=n3+1;    
end
end

end


%Edge Resize
imshow(Picture2);

if z == 1;
set(gcf, 'Units', 'Normalized', 'OuterPosition', [0 0 1 1]);
end
warning('off', 'images:initSize:adjustingMag');
hold on
title(['Questionnaire: ' num2str(z) '/' num2str(Nb_Files)])
drawnow;

%Well detected answers
for i = 1:[n2-1]; 
    rectangle('position', Feed_Mat(i,1:4), 'EdgeColor',Feed_Mat(i,5:7),'LineWidth',2,'Curvature',[0.8 0.8]);
end

%Missdetected answers
for i = 1:[n3-1];   
   rectangle('position', Feed_Mat2(i,1:4), 'EdgeColor',Feed_Mat2(i,5:7),'LineWidth',2,'Curvature',[0.8 0.8],'LineStyle','--');
end

hold on
%Question numbering
for i3 = 1:Nb_Question;
X = [Model(i3,4,1)-100];
Y = [Model(i3,2,1)]+5;
if X<1;
    X = 1;
end
text(X,Y, num2str(i3),'FontName', 'Arial', 'FontWeight', 'bold', 'FontSize', 14, 'Color', 'cyan')
end


%corrections of miss-detected answers
if n3>1;
    uiwait(helpdlg('Click on the right answer (if no answer exist, click in a  corner of the picture'));
    j2=1;
for j = 1:Nb_Question;
[Xv2 Yv2] = find(Feed_Mat2(:,8)==j);

if length(Yv2)>0;

imshow(Picture2);
set(gcf, 'Units', 'Normalized');
title(['Questionnaire: ' num2str(z) '/' num2str(Nb_Files)])
drawnow;

%Questions number
for i3 = 1:Nb_Question;
X = [Model(i3,4,1)-100];
Y = [Model(i3,2,1)]+5;
if X<1;
    X = 1;
end
text(X,Y, num2str(i3), 'FontName', 'Arial', 'FontWeight', 'bold', 'FontSize', 14, 'Color', 'cyan')
end
drawnow;

%Identification of missdetected answer
for r = 0:[nnz(Yv2)-1];
rectangle('position', Feed_Mat2([j2+r],1:4), 'EdgeColor',Feed_Mat2([j2+r],5:7),'LineWidth',2,'Curvature',[0.8 0.8],'LineStyle','--');
end
j2=j2+r+1;

% New answer selection
P1 = ginput(1);
t=1;
if P1(1,2)> [Model(j,2)*0.9] && P1(1,2)<[Model(j,3)*1.1];
while P1(1,1)>Model(j,[t+3]);
t = t+1;
end

Results2(j,2) = t-1;
Results2(j,4) = 1;
else
Results2(j,2) = nan;
Results2(j,4) = nan;   
end
end
end


imshow(Picture2);
warning('off', 'images:initSize:adjustingMag');
title(['Questionnaire: ' num2str(z) '/' num2str(Nb_Files)]);
drawnow;

for i = 1:Nb_Question
  if Results2(i,2)>0;
    Feed_Mat(i,2) = Model(i,2);
    Feed_Mat(i,1) = Model(i,[Results2(i,2)+3]);
    Feed_Mat(i,4) = Model(i,3)-Model(i,2);
    Feed_Mat(i,3) = Model(i,[Results2(i,2)+4])-Model(i,[Results2(i,2)+3]);
    Feed_Mat(i,5:7) = [0 1 0];
    rectangle('position', Feed_Mat(i,1:4), 'EdgeColor',Feed_Mat(i,5:7),'LineWidth',2,'Curvature',[0.8 0.8]);
  else
    Feed_Mat(i,2) = 0;
    Feed_Mat(i,1) = 0;
    Feed_Mat(i,4) = 0;
    Feed_Mat(i,3) = 0;
    Feed_Mat(i,5:7) = [0 1 0]; 
  end
  
X = [Model(i,4,1)-100];
Y = [Model(i,2,1)]+5;
if X<1;
    X = 1;
end
text(X,Y, num2str(i),'FontName', 'Arial', 'FontWeight', 'bold', 'FontSize', 14, 'Color', 'cyan');
end  


end

waitforbuttonpress;
Correction=1;
%Manual correction ask by user
while Correction>0;

CorrOK = false;
while CorrOK ~= true;
prompt = {'Are you agree with the answers decteted, if not, please indicate the number of the question you will change. Else click "ok"'};
dlg_title = 'Input';
num_lines = 1;
defaultans = {'0'};
answer = inputdlg(prompt,dlg_title,num_lines,defaultans);
Correction = str2num(answer{1});
if Correction<0 || Correction>Nb_Question;
uiwait(errordlg('Question selected must be a number comprise between 0 and the number of question in your questionnary'));
else
    CorrOK = true;
end
end

if Correction>0 && Correction < Nb_Question;
Feed_Correct(1,2) = Model (Correction,2);
Feed_Correct(1,1) = Model(Correction,4);
Feed_Correct(1,4) = Model(Correction,3)-Model(Correction,2);
Feed_Correct(1,3) = Model(Correction,[Nb_Choice+4])-Model(Correction,4);
Feed_Correct(1,5:7) = [1 0 0];

imshow(Picture2);
warning('off', 'images:initSize:adjustingMag');
title(['Questionnaire: ' num2str(z) '/' num2str(Nb_Files)]);
drawnow;

for i3 = 1:Nb_Question;
X = [Model(i3,4,1)-100];
Y = [Model(i3,2,1)]+5;
if X<1;
    X = 1;
end
a=text(X,Y, num2str(i3));
set(a, 'FontName', 'Arial', 'FontWeight', 'bold', 'FontSize', 14, 'Color', 'cyan');
end
rectangle('position', Feed_Correct(1,1:4), 'EdgeColor',Feed_Correct(1,5:7),'LineWidth',2,'Curvature',0.8,'LineStyle','--');
 
P1 = ginput(1);
t=1;
if P1(1,2)> [Model(Correction,2)*0.9] && P1(1,2)<[Model(Correction,3)*1.1];
while P1(1,1)>Model(Correction,[t+3]);
t = t+1;
end

Results2(Correction,2) = t-1;
Results2(Correction,4) = 1;
else
Results2(Correction,2) = nan;
Results2(Correction,4) = nan;   
end

  if Results2(Correction,2) > 0;
    Feed_Mat(Correction,2) = Model(Correction,2);
    Feed_Mat(Correction,1) = Model(Correction,[Results2(Correction,2)+3]);
    Feed_Mat(Correction,4) = Model(Correction,3)-Model(Correction,2);
    Feed_Mat(Correction,3) = Model(Correction,[Results2(Correction,2)+4])-Model(Correction,[Results2(Correction,2)+3]);
  else
    Feed_Mat(Correction,2) = 0;
    Feed_Mat(Correction,1) = 0;
    Feed_Mat(Correction,4) = 0;
    Feed_Mat(Correction,3) = 0;
  end
  
  
imshow(Picture2);
warning('off', 'images:initSize:adjustingMag');
title(['Questionnaire: ' num2str(z) '/' num2str(Nb_Files)]);
drawnow;

for i3 = 1:Nb_Question;
rectangle('position', Feed_Mat(i3,1:4), 'EdgeColor',Feed_Mat(i3,5:7),'LineWidth',2,'Curvature',[0.8 0.8]);
X = [Model(i3,4,1)-100];
Y = [Model(i3,2,1)]+5;
if X<1;
    X = 1;
end
a=text(X,Y, num2str(i3));
set(a, 'FontName', 'Arial', 'FontWeight', 'bold', 'FontSize', 14, 'Color', 'cyan');
end

waitforbuttonpress;

end
end
end
hold off


%Corection ofthe values atribuated to each answers


for r = 1:Nb_Question;
    if Results2(r,2)>0;
Results2(r,3) = Values (1, Results2(r,2));
    else
Results2(r,3) = nan;        
    end
end

%Inverted Scales
Nb_Inv = sum(Nb_Inv_Question > 0);
for r = 1:Nb_Inv;
Results2(Nb_Inv_Question(1,r),3) = (Values(1,Nb_Choice)+Values(1,1))-Results2(Nb_Inv_Question(1,r),2);
end

%%Increment datas in a Saving Matrix
Mat_Save(3:[Nb_Question+2],[s+1]:[s+3]) = Results2(1:[Nb_Question],2:4);

%Variables reinitialisation
s=s+3;
clear Feed_Mat
clear Feed_Mat2
clear Feed_Correct
Model(:,:,:) = Model2(:,:,:);
end

%Add the Question Number
Mat_Save(3:Nb_Question+2,1) = (1:1:Nb_Question)';
Title = cell(2,Nb_Files*3);
sub =1;
Title (2,1) = {'Question'};
Title (1,1) = {NaN};
for i = 2:3:[Nb_Files*3+1];
    Title (1,i) = {'Subject:'} ;
    Title(1,[i+1]) = {num2str(sub)};
    Title(1,[i+2]) = {NaN};
    Title(2,i:[i+2]) = {'Answers' 'Value of Answers' 'Nb of answers detected'};
    sub = sub+1;
end


% final matrix saving
if exist([Output_Folder '' MCQ_Model '-Analysis.xls'],'file'); %if file already exist with the same name
    List = dir([Output_Folder '' MCQ_Model '-Analysis' '*.xls']);
    disp('WARNING. Result file already exists with same name.')
    disp('It was saved as "Result_file_copy(n)".');
xlswrite([Output_Folder '' MCQ_Model '-Analysis_copy(' num2str(length(List)) ').xls'],Mat_Save);
xlswrite([Output_Folder '' MCQ_Model '-Analysis_copy(' num2str(length(List)) ').xls'],Title);
else      
xlswrite([Output_Folder '' MCQ_Model '-Analysis.xls'],Mat_Save);
xlswrite([Output_Folder '' MCQ_Model '-Analysis.xls'],Title);
end

%set(gcf, 'Visible', 'off');
close gcf

%%%%%%%%%%%%%%%%%%%%%%
%myicon = imread('Moi.jpg');
%h=msgbox('Operation Completed, Clovis you are so fucking good !!','Success','custom',myicon);
toc
```
