## Supplementary material for "Paper-and-pencil questionnaires analysis: a new automated technique to reduce analysis time and errors": LightQuest_script_package: LightQuest_Model.m

```
clear all
clc
tic

%%%%%%%%%%%%%%%%%%

prompt = {'Filename of the model (without ".jpg"):','Number of reference targets on questionnaire:','Number of Multiple Choice Question:'...
    'Number of choice by question:','Value of the answers (separated by space):','Number of the questions with inverted values (separated by space):'};
dlg_title = 'Input';
num_lines = 1;
defaultans = {'Model_QUEST','2','30', '5', '0 1 2 3 4', '0'};
%defaultans = {'example: POMS','example: 5', 'example: 5', 'example: 1 2 3 4 5', 'example: 6 15', 'example: 5'};
answer = inputdlg(prompt,dlg_title,num_lines,defaultans);


File_Name = answer{1};
Nb_Target = str2num(answer{2});
Nb_Question = str2double(answer(3,1));
Nb_Choice = str2num(answer{4});
Values = str2num(answer{5});
Nb_Inv_Question = str2num(answer{6});
Edge = 5;

%FOLDERS
Files_Folder = '../Files/'; % XML Folder
Model_Folder = '../Models/'; % folder containing the XLS result files

%%%%%%%%%%%%%%%%%%%%%%%%%%%%%

% Parameters testing
disp('>> TESTING PARAMETERS');
if ~exist(Files_Folder,'dir');
    uiwait(errordlg('Files folder is wrong. Please check Video_Folder.'));
elseif ~exist(Model_Folder);
    uiwait(errordlg('Model folder is wrong. Please check Output_Folder.'));
end
List_Files = dir([Files_Folder File_Name '.*']); %list of files in Files_Folder
if length(List_Files) < 1;
    uiwait(errordlg('Model file not found, please check if present or check the file name of the model'));   
elseif (Nb_Target<0 || Nb_Target>2);
    uiwait(helpdlg('The number of target selected is inferior to 2. Please draws 2 targets on your questionnaire to maximize analysis quality.'));
elseif (Nb_Question<1);
    uiwait(errordlg('The number Question selected is inferior to 1. Please change its value and run again script.'));
elseif (Nb_Choice<1);
    uiwait(errordlg('The number of choice by question is inferior to 1. Please change its value and run again script.'));
elseif (size(Values)~= Nb_Choice);
    uiwait(errordlg('The number of values attribuated to the different answers is not equal to the number of possible answers, please separate the values by a space (ex: 1 2 3)'));
elseif max(Nb_Inv_Question)> Nb_Question;
    uiwait(errordlg('At least one number of inverted question is over the number of question containing in this questionnary'));
else;
    disp('Parameters ok. Running script goes on.');
end

%%%%%%%%%%%%%%%%%%%%%%%%%%%%%

Picture = imread([Files_Folder '' File_Name '.jpg']);


[Ymax Xmax] = size(Picture);
Model = zeros (Nb_Question,50,2);
Screen = get(0, 'ScreenSize');
if Nb_Target>0;
% Manual picking of target references(example : Rodent cage rectangle)
imshow(Picture); warning('off', 'images:initSize:adjustingMag'); axis on;
set(gcf, 'Position', Screen);
hold on
uiwait(msgbox({'Select the area contening the first black rectangle of reference' '(top left corner to bottom right, keep mouse button pressed)'}));
Ar =1;
C=0;
while Ar <= Nb_Target;

xlabel('X-Axis (pixels)');
ylabel('Y-Axis (pixels)');
title('Definition of reference targets');
if Ar == 2 && C ==0;
uiwait(msgbox({'Select the area contening the second black rectangle of reference' '(top left corner to bottom right, keep mouse button pressed)'}));
end


AreaOK = false;
while AreaOK ~= true;
    set(gcf,'Units','normalized')
    k = waitforbuttonpress;
    P1 = get(gca,'CurrentPoint');
    rect_pos = rbbox;
    if Ar == 1
    Rec1 = annotation('rectangle',rect_pos,'Color','black', 'LineStyle', '-.','Tag', 'Rec1');
    else
    Rec2 = annotation('rectangle',rect_pos,'Color','black', 'LineStyle', '-.','Tag', 'Rec2'); 
    end
    drawnow;
    P2 = get(gca,'CurrentPoint');
    X1 = round(P1(1,1))-20;
    X2 = round(P2(1,1))+20;
    Y1 = round(P1(1,2))-20;
    Y2 = round(P2(1,2))+20;
    if X1 < 0; X1= 1; end
    if Y1 < 0; Y1 = 1; end
    if X2 > Xmax; X2 = Xmax; end
    if Y2> Ymax; Y2 = Ymax; end
    
    if (X2 < X1) || (Y2 < Y1) || (X1 <= 0) || (X2 <= 0) || (Y1 <= 0) || (Y2 <= 0) || [(X1-X2)*(Y1-Y2)]<10000 ;
        uiwait(errordlg({'Incorrect area. Please enter new target area' '(top left corner to bottom right, keep mouse button pressed)'}));
        AreaOK = false;
        if Ar ==1;   
        delete(findall(gcf,'Tag','Rec1'));
        else
        delete(findall(gcf,'Tag','Rec2'));
        end
    else;
        AreaOK = true;
    end  
    
end 

Pic = Picture(Y1:Y2,X1:X2);
Pic = imcomplement(im2bw(Pic,0.9));
Pic = bwareaopen(Pic,300);
bw = bwlabel(Pic, 4);
stats1 = regionprops(bw, 'Centroid','Area');


if length(stats1)>0;
Model([6+Ar],1,2)= stats1(1).Area;
Model([6+Ar],2:3,2)= stats1(1).Centroid;
Model([6+Ar],2,2)=Model([6+Ar],2,2)+X1;
Model([6+Ar],3,2)=Model([6+Ar],3,2)+Y1;
Model([6+Ar],4,2)=Y1;
Model([6+Ar],5,2)=Y2;
Model([6+Ar],6,2)=X1;
Model([6+Ar],7,2)=X2;
Ar =Ar+1;
else
uiwait(errordlg({'No target detected. Please select again' '(top left corner to bottom right, keep mouse button pressed)'})); 
if Ar ==1;   
delete(findall(gcf,'Tag','Rec1'));
else
C = 1;
delete(findall(gcf,'Tag','Rec2'));
end
end


end

hold off

%Rotation of the Picture
if Nb_Target>1;
Rot_cor(1,1) = abs(Model(7,2,2)-Model(8,2,2));
Rot_cor(2,1) = abs(Model(7,3,2)-Model(8,3,2));
S_Pic = size(Picture);
Picture = imrotate(Picture,round([atand(Rot_cor(2,1)/Rot_cor(1,1))],3));
Mrot = ~imrotate(true(S_Pic),[atand(Rot_cor(2,1)/Rot_cor(1,1))]);
Picture(Mrot&~imclearborder(Mrot)) = 255;
[Ymax Xmax] = size(Picture);
end

%Detection of the new centroid of the first target
Pic2 = Picture(Model(7,4,2):Model(7,5,2),Model(7,6,2):Model(7,7,2));
Pic2 = imcomplement(im2bw(Pic2,0.9));
Pic2 = bwareaopen(Pic2,300);
bw = bwlabel(Pic2, 4);
stats2 = regionprops(bw, 'Centroid');
Model(9,2:3,2)= stats2(1).Centroid;
Model(9,2,2)=Model(9,2,2)+ Model(7,6,2);
Model(9,3,2)=Model(9,3,2)+ Model(7,4,2);
end

%transformation of Picture in binary
Picture = im2bw(Picture,0.9);

% Manual picking of Questions area
imshow(Picture); warning('off', 'images:initSize:adjustingMag'); axis on;
%set(gcf, 'Position', Screen);
if Nb_Target == 0;
end
xlabel('X-Axis (pixels)');
ylabel('Y-Axis (pixels)');
title('Definition of questions area');
uiwait(msgbox({'Select the Questions area' '(top left corner to bottom right, keep mouse button pressed)'}));
AreaOK = false;
while AreaOK ~= true;
    set(gcf,'Units','normalized')
    k = waitforbuttonpress;
    P1 = get(gca,'CurrentPoint');
    rect_pos = rbbox;
    Rec1 = annotation('rectangle',rect_pos,'Color','cyan');
    P2 = get(gca,'CurrentPoint');
    X1 = round(P1(1,1));
    X2 = round(P2(1,1));
    Y1 = round(P1(1,2));
    Y2 = round(P2(1,2));
    if X1 < 0; X1= 1; end
    if Y1 < 0; Y1 = 1; end
    if X2 > Xmax; X2 = Xmax; end
    if Y2> Ymax; Y2 = Ymax; end
    if (X2 < X1) || (Y2 < Y1) || (X1 <= 0) || (X2 <= 0) || (Y1 <= 0) || (Y2 <= 0)|| [(X1-X2)*(Y1-Y2)]<10000;
        uiwait(errordlg({'Incorrect area. Please enter new Questions area' '(top left corner to bottom right, keep mouse button pressed)'}));
        AreaOK = false;
    else;
        AreaOK = true;
    end
    %delete(Rec1);
end 
clf

Picture2 = Picture(Y1:Y2,X1:X2);
[Ymax2 Xmax2] = size(Picture2);

Model(10,4,2)=Y1;
Model(10,5,2)=Y2;
Model(10,6,2)=X1;
Model(10,7,2)=X2;


% Manual picking of target area for each question 
Answers_Coordinates = zeros (Nb_Question,5);
i=1;
imshow(Picture2); warning('off', 'images:initSize:adjustingMag');axis on;hold on; grid on; grid minor;
set(gcf, 'Units', 'Normalized', 'OuterPosition', [0 0 1 1]);
uiwait(msgbox({'Select all the answer area of each question in one rectangle' '(top left corner to bottom right, keep mouse button pressed)'}));
for i=1:Nb_Question;
xlabel('X-Axis (pixels)');
ylabel('Y-Axis (pixels)');
title(['Definition of Question area:' num2str(i)]);
% Test if correct target area
AreaOK = false;
while AreaOK ~= true;
    
    set(gcf,'Units','normalized')
    k = waitforbuttonpress;
    P1 = get(gca,'CurrentPoint');
    rect_pos = rbbox;
    Rec1 = annotation('rectangle',rect_pos,'Color','red', 'LineStyle','-.');
    P2 = get(gca,'CurrentPoint');
    Answers_Coordinates(i,1) = round(P1(1,1));
    Answers_Coordinates(i,2) = round(P2(1,1));
    Answers_Coordinates(i,3) = round(P1(1,2));
    Answers_Coordinates(i,4) = round(P2(1,2));
    if Answers_Coordinates(i,1) < 0; Answers_Coordinates(i,1)= 1; end
    if Answers_Coordinates(i,3) < 0; Answers_Coordinates(i,3) = 1; end
    if Answers_Coordinates(i,2) > Xmax2; Answers_Coordinates(i,2) = Xmax2; end
    if Answers_Coordinates(i,4)> Ymax2; Answers_Coordinates(i,4) = Ymax2; end
    if (Answers_Coordinates(i,2) < Answers_Coordinates(i,1)) || (Answers_Coordinates(i,4) < Answers_Coordinates(i,3)) | ((Answers_Coordinates(i,2)-Answers_Coordinates(i,1))*(Answers_Coordinates(i,4)-(Answers_Coordinates(i,3))))<10000 ;
        uiwait(errordlg({'Incorrect area. Please enter new Question area' '(top left corner to bottom right, keep mouse button pressed)'}));
        AreaOK = false;
    else;
        AreaOK = true;
    end
end 

end

%Question Numerotations
for i = 1:Nb_Question;
 X3 = [Answers_Coordinates(i,1)-100];
 Y3 = [Answers_Coordinates(i,3)]+5;
  if X3<1;
    X3 = 1;
  end
hold on
a=text(X3,Y3, num2str(i));
set(a, 'FontName', 'Arial', 'FontWeight', 'bold', 'FontSize', 16, 'Color', 'cyan');
end

%Increment of value in Matrix save
for i = 1:Nb_Question;
    R_Size = round(((Answers_Coordinates(i,2)-Answers_Coordinates(i,1))/Nb_Choice)-0.5);

    Model(i,2,1) = Answers_Coordinates(i,3);
    Model(i,3,1) = Answers_Coordinates(i,4);
    Model(i,4,1) = Answers_Coordinates(i,1);
    
  for q =1:Nb_Choice
    Model(i,[q+4],1) = [Model(i,4,1)+[q*R_Size]];
    Model(i,[q+4+Nb_Choice],1) = nnz(~(Picture2([Model(i,2,1)]:[Model(i,3,1)],[Model(i,[q+3],1)+Edge]:[Model(i,[q+4],1)-Edge])));
   
  end
end

hold off

%Increment of variables in the saving matrix

Model(2,1,2) = Nb_Question;
Model(3,1,2) = Nb_Choice;
Model(4,1:Nb_Choice,2) = Values;
S_NbIV = nnz(Nb_Inv_Question);
Model(5,1:S_NbIV,2) = Nb_Inv_Question;
Model(6,1,2) = Edge;
Model(7,1,2) = Nb_Target;


%%%%%%%%%%%%%%%%%%%%%%%%%%%%%%%%%%%%%%%%%%%%%
%Correction of the model
Correction = 1;
Feed_Correct = zeros(1,7);
%rectangles coordinates calculation
for n3 = 1:Nb_Question;
Feed_Mat(n3,2) = Model(n3,2);
Feed_Mat(n3,1) = Model(n3,4);
Feed_Mat(n3,4) = Model(n3,3)-Model(n3,2);
Feed_Mat(n3,3) = Model(n3,[Nb_Choice+4])-Model(n3,4);
end 

while Correction>0;
CorrOK = false;
while CorrOK ~= true;
prompt = {'Are you agree with the areas decteted, if not, please indicate the number of the question you will change and re-select the area you want. Else click "ok"'};
dlg_title = 'Input';
num_lines = 1;
defaultans = {'0'};
answer = inputdlg(prompt,dlg_title,num_lines,defaultans);
Correction = str2num(answer{1});

if Correction<0 || Correction>Nb_Question;
uiwait(errordlg('Question selected must be a number comprise between 0 and the number of question in your questionnary'));
else
    CorrOK = true;
end
end


%If correction is wanted
if Correction>0;
clf
imshow(Picture2); hold on;
set(gcf, 'Units', 'Normalized', 'OuterPosition', [0 0 1 1]);hold on; axis on; grid on; grid minor;
rectangle('position', Feed_Mat(Correction,1:4), 'EdgeColor','red','LineWidth',1, 'LineStyle', '-.');

xlabel('X-Axis (pixels)');
ylabel('Y-Axis (pixels)');
title(['Definition of Question area:' num2str(Correction)]);

%selection of the new question area
AreaOK = false;
while AreaOK ~= true;
    
    set(gcf,'Units','normalized')
    k = waitforbuttonpress;
    P1 = get(gca,'CurrentPoint');
    rect_pos = rbbox;
    Rec1 = annotation('rectangle',rect_pos,'Color','red');
    P2 = get(gca,'CurrentPoint');
    Answers_Coordinates(1,1) = round(P1(1,1));
    Answers_Coordinates(1,2) = round(P2(1,1));
    Answers_Coordinates(1,3) = round(P1(1,2));
    Answers_Coordinates(1,4) = round(P2(1,2));
    if Answers_Coordinates(1,1) < 0; Answers_Coordinates(1,1)= 1; end
    if Answers_Coordinates(1,3) < 0; Answers_Coordinates(1,3) = 1; end
    if Answers_Coordinates(1,2) > Xmax2; Answers_Coordinates(1,2) = Xmax2; end
    if Answers_Coordinates(1,4)> Ymax2; Answers_Coordinates(1,4) = Ymax2; end
    if (Answers_Coordinates(1,2) < Answers_Coordinates(1,1)) | (Answers_Coordinates(1,4) < Answers_Coordinates(1,3)) | ((Answers_Coordinates(1,2)-Answers_Coordinates(1,1))*(Answers_Coordinates(1,4)-(Answers_Coordinates(1,3))))<10000 ;
        uiwait(errordlg({'Incorrect area. Please enter new Question area' '(top left corner to bottom right, keep mouse button pressed)'}));
        AreaOK = false;
    else;
        AreaOK = true;
    end
end 
hold off

%correction of coordinates
Answers_Coordinates(1,1) = [Answers_Coordinates(1,1)];
Answers_Coordinates(1,2) = [Answers_Coordinates(1,2)];
Answers_Coordinates(1,3) = [Answers_Coordinates(1,3)];
Answers_Coordinates(1,4) = [Answers_Coordinates(1,4)];

%Increment of new answer coordinates
    Model(Correction,2,1) = Answers_Coordinates(1,3);
    Model(Correction,3,1) = Answers_Coordinates(1,4);
    Model(Correction,4,1) = Answers_Coordinates(1,1);
    
 R_Size2 = round(((Answers_Coordinates(Correction,2)-Answers_Coordinates(Correction,1))/Nb_Choice)-0.5);   
  for q =1:Nb_Choice
    Model(Correction,[q+4],1) = [Model(Correction,4,1)+[q*R_Size2]];
    Model(Correction,[q+4+Nb_Choice],1) = nnz(~(Picture([Model(Correction,2,1)]:[Model(Correction,3,1)],[Model(Correction,[q+3],1)+Edge]:[Model(Correction,[q+4],1)-Edge]))); 
  end
  
Feed_Mat(Correction,2) = Model(Correction,2);
Feed_Mat(Correction,1) = Model(Correction,4);
Feed_Mat(Correction,4) = Model(Correction,3) - Model(Correction,2);
Feed_Mat(Correction,3) = Model(Correction,[Nb_Choice+4]) - Model(Correction,4);
end


%Visual controlof the answer's areas
clf
imshow(Picture2); hold on;
set(gcf, 'Units', 'Normalized', 'OuterPosition', [0 0 1 1]);
for i = 1: Nb_Question
rectangle('position', Feed_Mat(i,1:4), 'EdgeColor','red','LineWidth',1);
X = [Model(i,4,1)-100];
Y = [Model(i,2,1)]+5;
if X < 1;
   X = 1;
end
a=text(X,Y, num2str(i));
set(a, 'FontName', 'Arial', 'FontWeight', 'bold', 'FontSize', 10, 'Color', 'cyan');
end
hold off
end

 % Result export
    Savefile=[Model_Folder '' File_Name '.mat'];
    List = dir([Model_Folder '' File_Name '*.mat']);
    if exist(Savefile,'file'); %Changes the name of the export file if it already exist in the folder 
        disp('WARNING. Matrix file already exists with same name. Saving as Matrix_file_copy.');
        save([Model_Folder '' File_Name '_copy(' num2str(length(List)) ').mat'],'Model');
    else;
        disp('Saving Matrix_file');
        save(Savefile,'Model');
    end
  close gcf
  close all
  toc
```
