## Supplementary material for "Paper-and-pencil questionnaires analysis: a new automated technique to reduce analysis time and errors": LightQuest_script_package: LICENSE

                                 Apache License
                           Version 2.0, January 2004
                        http://www.apache.org/licenses/

   TERMS AND CONDITIONS FOR USE, REPRODUCTION, AND DISTRIBUTION

   1. Definitions.

      "License" shall mean the terms and conditions for use, reproduction,
      and distribution as defined by Sections 1 through 9 of this document.

      "Licensor" shall mean the copyright owner or entity authorized by
      the copyright owner that is granting the License.

      "Legal Entity" shall mean the union of the acting entity and all
      other entities that control, are controlled by, or are under common
      control with that entity. For the purposes of this definition,
      "control" means (i) the power, direct or indirect, to cause the
      direction or management of such entity, whether by contract or
      otherwise, or (ii) ownership of fifty percent (50%) or more of the
      outstanding shares, or (iii) beneficial ownership of such entity.

      "You" (or "Your") shall mean an individual or Legal Entity
      exercising permissions granted by this License.

      "Source" form shall mean the preferred form for making modifications,
      including but not limited to software source code, documentation
      source, and configuration files.

      "Object" form shall mean any form resulting from mechanical
      transformation or translation of a Source form, including but
      not limited to compiled object code, generated documentation,
      and conversions to other media types.

      "Work" shall mean the work of authorship, whether in Source or
      Object form, made available under the License, as indicated by a
      copyright notice that is included in or attached to the work
      (an example is provided in the Appendix below).

      "Derivative Works" shall mean any work, whether in Source or Object
      form, that is based on (or derived from) the Work and for which the
      editorial revisions, annotations, elaborations, or other modifications
      represent, as a whole, an original work of authorship. For the purposes
      of this License, Derivative Works shall not include works that remain
      separable from, or merely link (or bind by name) to the interfaces of,
      the Work and Derivative Works thereof.

      "Contribution" shall mean any work of authorship, including
      the original version of the Work and any modifications or additions
      to that Work or Derivative Works thereof, that is intentionally
      submitted to Licensor for inclusion in the Work by the copyright owner
      or by an individual or Legal Entity authorized to submit on behalf of
      the copyright owner. For the purposes of this definition, "submitted"
      means any form of electronic, verbal, or written communication sent
      to the Licensor or its representatives, including but not limited to
      communication on electronic mailing lists, source code control systems,
      and issue tracking systems that are managed by, or on behalf of, the
      Licensor for the purpose of discussing and improving the Work, but
      excluding communication that is conspicuously marked or otherwise
      designated in writing by the copyright owner as "Not a Contribution."

      "Contributor" shall mean Licensor and any individual or Legal Entity
      on behalf of whom a Contribution has been received by Licensor and
      subsequently incorporated within the Work.

   2. Grant of Copyright License. Subject to the terms and conditions of
      this License, each Contributor hereby grants to You a perpetual,
      worldwide, non-exclusive, no-charge, royalty-free, irrevocable
      copyright license to reproduce, prepare Derivative Works of,
      publicly display, publicly perform, sublicense, and distribute the
      Work and such Derivative Works in Source or Object form.

   3. Grant of Patent License. Subject to the terms and conditions of
      this License, each Contributor hereby grants to You a perpetual,
      worldwide, non-exclusive, no-charge, royalty-free, irrevocable
      (except as stated in this section) patent license to make, have made,
      use, offer to sell, sell, import, and otherwise transfer the Work,
      where such license applies only to those patent claims licensable
      by such Contributor that are necessarily infringed by their
      Contribution(s) alone or by combination of their Contribution(s)
      with the Work to which such Contribution(s) was submitted. If You
      institute patent litigation against any entity (including a
      cross-claim or counterclaim in a lawsuit) alleging that the Work
      or a Contribution incorporated within the Work constitutes direct
      or contributory patent infringement, then any patent licenses
      granted to You under this License for that Work shall terminate
      as of the date such litigation is filed.

   4. Redistribution. You may reproduce and distribute copies of the
      Work or Derivative Works thereof in any medium, with or without
      modifications, and in Source or Object form, provided that You
      meet the following conditions:

      (a) You must give any other recipients of the Work or
          Derivative Works a copy of this License; and

      (b) You must cause any modified files to carry prominent notices
          stating that You changed the files; and

      (c) You must retain, in the Source form of any Derivative Works
          that You distribute, all copyright, patent, trademark, and
          attribution notices from the Source form of the Work,
          excluding those notices that do not pertain to any part of
          the Derivative Works; and

      (d) If the Work includes a "NOTICE" text file as part of its
          distribution, then any Derivative Works that You distribute must
          include a readable copy of the attribution notices contained
          within such NOTICE file, excluding those notices that do not
          pertain to any part of the Derivative Works, in at least one
          of the following places: within a NOTICE text file distributed
          as part of the Derivative Works; within the Source form or
          documentation, if provided along with the Derivative Works; or,
          within a display generated by the Derivative Works, if and
          wherever such third-party notices normally appear. The contents
          of the NOTICE file are for informational purposes only and
          do not modify the License. You may add Your own attribution
          notices within Derivative Works that You distribute, alongside
          or as an addendum to the NOTICE text from the Work, provided
          that such additional attribution notices cannot be construed
          as modifying the License.

      You may add Your own copyright statement to Your modifications and
      may provide additional or different license terms and conditions
      for use, reproduction, or distribution of Your modifications, or
      for any such Derivative Works as a whole, provided Your use,
      reproduction, and distribution of the Work otherwise complies with
      the conditions stated in this License.

   5. Submission of Contributions. Unless You explicitly state otherwise,
      any Contribution intentionally submitted for inclusion in the Work
      by You to the Licensor shall be under the terms and conditions of
      this License, without any additional terms or conditions.
      Notwithstanding the above, nothing herein shall supersede or modify
      the terms of any separate license agreement you may have executed
      with Licensor regarding such Contributions.

   6. Trademarks. This License does not grant permission to use the trade
      names, trademarks, service marks, or product names of the Licensor,
      except as required for reasonable and customary use in describing the
      origin of the Work and reproducing the content of the NOTICE file.

   7. Disclaimer of Warranty. Unless required by applicable law or
      agreed to in writing, Licensor provides the Work (and each
      Contributor provides its Contributions) on an "AS IS" BASIS,
      WITHOUT WARRANTIES OR CONDITIONS OF ANY KIND, either express or
      implied, including, without limitation, any warranties or conditions
      of TITLE, NON-INFRINGEMENT, MERCHANTABILITY, or FITNESS FOR A
      PARTICULAR PURPOSE. You are solely responsible for determining the
      appropriateness of using or redistributing the Work and assume any
      risks associated with Your exercise of permissions under this License.

   8. Limitation of Liability. In no event and under no legal theory,
      whether in tort (including negligence), contract, or otherwise,
      unless required by applicable law (such as deliberate and grossly
      negligent acts) or agreed to in writing, shall any Contributor be
      liable to You for damages, including any direct, indirect, special,
      incidental, or consequential damages of any character arising as a
      result of this License or out of the use or inability to use the
      Work (including but not limited to damages for loss of goodwill,
      work stoppage, computer failure or malfunction, or any and all
      other commercial damages or losses), even if such Contributor
      has been advised of the possibility of such damages.

   9. Accepting Warranty or Additional Liability. While redistributing
      the Work or Derivative Works thereof, You may choose to offer,
      and charge a fee for, acceptance of support, warranty, indemnity,
      or other liability obligations and/or rights consistent with this
      License. However, in accepting such obligations, You may act only
      on Your own behalf and on Your sole responsibility, not on behalf
      of any other Contributor, and only if You agree to indemnify,
      defend, and hold each Contributor harmless for any liability
      incurred by, or claims asserted against, such Contributor by reason
      of your accepting any such warranty or additional liability.

   END OF TERMS AND CONDITIONS

   APPENDIX: How to apply the Apache License to your work.

      To apply the Apache License to your work, attach the following
      boilerplate notice, with the fields enclosed by brackets "[]"
      replaced with your own identifying information. (Don't include
      the brackets!)  The text should be enclosed in the appropriate
      comment syntax for the file format. We also recommend that a
      file or class name and description of purpose be included on the
      same "printed page" as the copyright notice for easier
      identification within third-party archives.

   Copyright [yyyy] [name of copyright owner]

   Licensed under the Apache License, Version 2.0 (the "License");
   you may not use this file except in compliance with the License.
   You may obtain a copy of the License at

       http://www.apache.org/licenses/LICENSE-2.0

   Unless required by applicable law or agreed to in writing, software
   distributed under the License is distributed on an "AS IS" BASIS,
   WITHOUT WARRANTIES OR CONDITIONS OF ANY KIND, either express or implied.
   See the License for the specific language governing permissions and
   limitations under the License.


APACHE POI SUBCOMPONENTS:

Apache POI includes subcomponents with separate copyright notices and
license terms. Your use of these subcomponents is subject to the terms
and conditions of the following licenses:


Office Open XML schemas (ooxml-schemas-1.1.jar)

    The Office Open XML schema definitions used by Apache POI are
    a part of the Office Open XML ECMA Specification (ECMA-376, [1]).
    As defined in section 9.4 of the ECMA bylaws [2], this specification
    is available to all interested parties without restriction:

        9.4 All documents when approved shall be made available to
            all interested parties without restriction.

    Furthermore, both Microsoft and Adobe have granted patent licenses
    to this work [3,4,5].

    [1] http://www.ecma-international.org/publications/standards/Ecma-376.htm
    [2] http://www.ecma-international.org/memento/Ecmabylaws.htm
    [3] http://www.microsoft.com/interop/osp/
    [4] http://www.ecma-international.org/publications/files/ECMA-ST/Ecma%20PATENT/ECMA-376%20Edition%201%20Microsoft%20Patent%20Declaration.pdf
    [5] http://www.ecma-international.org/publications/files/ECMA-ST/Ecma%20PATENT/ga-2006-191.pdf


DOM4J library (dom4j-1.6.1.jar)

    Copyright 2001-2005 (C) MetaStuff, Ltd. All Rights Reserved.

    Redistribution and use of this software and associated documentation
    ("Software"), with or without modification, are permitted provided
    that the following conditions are met:

    1. Redistributions of source code must retain copyright
       statements and notices.  Redistributions must also contain a
       copy of this document.

    2. Redistributions in binary form must reproduce the
       above copyright notice, this list of conditions and the
       following disclaimer in the documentation and/or other
       materials provided with the distribution.

    3. The name "DOM4J" must not be used to endorse or promote
       products derived from this Software without prior written
       permission of MetaStuff, Ltd.  For written permission,
       please contact.

    4. Products derived from this Software may not be called "DOM4J"
       nor may "DOM4J" appear in their names without prior written
       permission of MetaStuff, Ltd. DOM4J is a registered
       trademark of MetaStuff, Ltd.

    5. Due credit should be given to the DOM4J Project - 
       http://www.dom4j.org
 
    THIS SOFTWARE IS PROVIDED BY METASTUFF, LTD. AND CONTRIBUTORS
    ``AS IS'' AND ANY EXPRESSED OR IMPLIED WARRANTIES, INCLUDING, BUT
    NOT LIMITED TO, THE IMPLIED WARRANTIES OF MERCHANTABILITY AND
    FITNESS FOR A PARTICULAR PURPOSE ARE DISCLAIMED.  IN NO EVENT SHALL
    METASTUFF, LTD. OR ITS CONTRIBUTORS BE LIABLE FOR ANY DIRECT,
    INDIRECT, INCIDENTAL, SPECIAL, EXEMPLARY, OR CONSEQUENTIAL DAMAGES
    (INCLUDING, BUT NOT LIMITED TO, PROCUREMENT OF SUBSTITUTE GOODS OR
    SERVICES; LOSS OF USE, DATA, OR PROFITS; OR BUSINESS INTERRUPTION)
    HOWEVER CAUSED AND ON ANY THEORY OF LIABILITY, WHETHER IN CONTRACT,
    STRICT LIABILITY, OR TORT (INCLUDING NEGLIGENCE OR OTHERWISE)
    ARISING IN ANY WAY OUT OF THE USE OF THIS SOFTWARE, EVEN IF ADVISED
    OF THE POSSIBILITY OF SUCH DAMAGE.


JUnit test library (junit-3.8.1.jar)

    Common Public License - v 1.0

    THE ACCOMPANYING PROGRAM IS PROVIDED UNDER THE TERMS OF THIS COMMON
    PUBLIC LICENSE ("AGREEMENT"). ANY USE, REPRODUCTION OR DISTRIBUTION
    OF THE PROGRAM CONSTITUTES RECIPIENT'S ACCEPTANCE OF THIS AGREEMENT.

    1. DEFINITIONS

    "Contribution" means:

    a) in the case of the initial Contributor, the initial code and
       documentation distributed under this Agreement, and

    b) in the case of each subsequent Contributor:

       i)  changes to the Program, and

       ii) additions to the Program;

       where such changes and/or additions to the Program originate from
       and are distributed by that particular Contributor. A Contribution
       'originates' from a Contributor if it was added to the Program by
       such Contributor itself or anyone acting on such Contributor's behalf.
       Contributions do not include additions to the Program which: (i) are
       separate modules of software distributed in conjunction with the
       Program under their own license agreement, and (ii) are not derivative
       works of the Program.

    "Contributor" means any person or entity that distributes the Program.

    "Licensed Patents " mean patent claims licensable by a Contributor which
    are necessarily infringed by the use or sale of its Contribution alone
    or when combined with the Program.

    "Program" means the Contributions distributed in accordance with this
    Agreement.

    "Recipient" means anyone who receives the Program under this Agreement,
    including all Contributors.

    2. GRANT OF RIGHTS

    a) Subject to the terms of this Agreement, each Contributor hereby grants
       Recipient a non-exclusive, worldwide, royalty-free copyright license
       to reproduce, prepare derivative works of, publicly display, publicly
       perform, distribute and sublicense the Contribution of such
       Contributor, if any, and such derivative works, in source code and
       object code form.

    b) Subject to the terms of this Agreement, each Contributor hereby grants
       Recipient a non-exclusive, worldwide, royalty-free patent license under
       Licensed Patents to make, use, sell, offer to sell, import and
       otherwise transfer the Contribution of such Contributor, if any, in
       source code and object code form. This patent license shall apply to
       the combination of the Contribution and the Program if, at the time
       the Contribution is added by the Contributor, such addition of the
       Contribution causes such combination to be covered by the Licensed
       Patents. The patent license shall not apply to any other combinations
       which include the Contribution. No hardware per se is licensed
       hereunder.

    c) Recipient understands that although each Contributor grants the
       licenses to its Contributions set forth herein, no assurances are
       provided by any Contributor that the Program does not infringe the
       patent or other intellectual property rights of any other entity.
       Each Contributor disclaims any liability to Recipient for claims
       brought by any other entity based on infringement of intellectual
       property rights or otherwise. As a condition to exercising the rights
       and licenses granted hereunder, each Recipient hereby assumes sole
       responsibility to secure any other intellectual property rights
       needed, if any. For example, if a third party patent license is
       required to allow Recipient to distribute the Program, it is
       Recipient's responsibility to acquire that license before
       distributing the Program.

    d) Each Contributor represents that to its knowledge it has sufficient
       copyright rights in its Contribution, if any, to grant the copyright
       license set forth in this Agreement.

    3. REQUIREMENTS

    A Contributor may choose to distribute the Program in object code form
    under its own license agreement, provided that:

    a) it complies with the terms and conditions of this Agreement; and

    b) its license agreement:

       i)   effectively disclaims on behalf of all Contributors all warranties
            and conditions, express and implied, including warranties or
            conditions of title and non-infringement, and implied warranties
            or conditions of merchantability and fitness for a particular
            purpose;

       ii)  effectively excludes on behalf of all Contributors all liability
            for damages, including direct, indirect, special, incidental and
            consequential damages, such as lost profits;

       iii) states that any provisions which differ from this Agreement are
            offered by that Contributor alone and not by any other party; and

       iv)  states that source code for the Program is available from such
            Contributor, and informs licensees how to obtain it in a
            reasonable manner on or through a medium customarily used for
            software exchange.

    When the Program is made available in source code form:

    a) it must be made available under this Agreement; and

    b) a copy of this Agreement must be included with each copy of
       the Program.

    Contributors may not remove or alter any copyright notices contained
    within the Program.

    Each Contributor must identify itself as the originator of its
    Contribution, if any, in a manner that reasonably allows subsequent
    Recipients to identify the originator of the Contribution.

    4. COMMERCIAL DISTRIBUTION

    Commercial distributors of software may accept certain responsibilities
    with respect to end users, business partners and the like. While this
    license is intended to facilitate the commercial use of the Program,
    the Contributor who includes the Program in a commercial product offering
    should do so in a manner which does not create potential liability for
    other Contributors. Therefore, if a Contributor includes the Program
    in a commercial product offering, such Contributor ("Commercial
    Contributor") hereby agrees to defend and indemnify every other
    Contributor ("Indemnified Contributor") against any losses, damages
    and costs (collectively "Losses") arising from claims, lawsuits and
    other legal actions brought by a third party against the Indemnified
    Contributor to the extent caused by the acts or omissions of such
    Commercial Contributor in connection with its distribution of the
    Program in a commercial product offering. The obligations in this
    section do not apply to any claims or Losses relating to any actual
    or alleged intellectual property infringement. In order to qualify,
    an Indemnified Contributor must: a) promptly notify the Commercial
    Contributor in writing of such claim, and b) allow the Commercial
    Contributor to control, and cooperate with the Commercial Contributor
    in, the defense and any related settlement negotiations. The Indemnified
    Contributor may participate in any such claim at its own expense.

    For example, a Contributor might include the Program in a commercial
    product offering, Product X. That Contributor is then a Commercial
    Contributor. If that Commercial Contributor then makes performance
    claims, or offers warranties related to Product X, those performance
    claims and warranties are such Commercial Contributor's responsibility
    alone. Under this section, the Commercial Contributor would have to
    defend claims against the other Contributors related to those
    performance claims and warranties, and if a court requires any other
    Contributor to pay any damages as a result, the Commercial Contributor
    must pay those damages.

    5. NO WARRANTY

    EXCEPT AS EXPRESSLY SET FORTH IN THIS AGREEMENT, THE PROGRAM IS PROVIDED
    ON AN "AS IS" BASIS, WITHOUT WARRANTIES OR CONDITIONS OF ANY KIND, EITHER
    EXPRESS OR IMPLIED INCLUDING, WITHOUT LIMITATION, ANY WARRANTIES OR
    CONDITIONS OF TITLE, NON-INFRINGEMENT, MERCHANTABILITY OR FITNESS FOR
    A PARTICULAR PURPOSE. Each Recipient is solely responsible for
    determining the appropriateness of using and distributing the Program
    and assumes all risks associated with its exercise of rights under this
    Agreement, including but not limited to the risks and costs of program
    errors, compliance with applicable laws, damage to or loss of data,
    programs or equipment, and unavailability or interruption of operations.

    6. DISCLAIMER OF LIABILITY

    EXCEPT AS EXPRESSLY SET FORTH IN THIS AGREEMENT, NEITHER RECIPIENT NOR
    ANY CONTRIBUTORS SHALL HAVE ANY LIABILITY FOR ANY DIRECT, INDIRECT,
    INCIDENTAL, SPECIAL, EXEMPLARY, OR CONSEQUENTIAL DAMAGES (INCLUDING
    WITHOUT LIMITATION LOST PROFITS), HOWEVER CAUSED AND ON ANY THEORY OF
    LIABILITY, WHETHER IN CONTRACT, STRICT LIABILITY, OR TORT (INCLUDING
    NEGLIGENCE OR OTHERWISE) ARISING IN ANY WAY OUT OF THE USE OR
    DISTRIBUTION OF THE PROGRAM OR THE EXERCISE OF ANY RIGHTS GRANTED
    HEREUNDER, EVEN IF ADVISED OF THE POSSIBILITY OF SUCH DAMAGES.

    7. GENERAL

    If any provision of this Agreement is invalid or unenforceable under
    applicable law, it shall not affect the validity or enforceability of
    the remainder of the terms of this Agreement, and without further
    action by the parties hereto, such provision shall be reformed to the
    minimum extent necessary to make such provision valid and enforceable.

    If Recipient institutes patent litigation against a Contributor with
    respect to a patent applicable to software (including a cross-claim or
    counterclaim in a lawsuit), then any patent licenses granted by that
    Contributor to such Recipient under this Agreement shall terminate as of
    the date such litigation is filed. In addition, if Recipient institutes
    patent litigation against any entity (including a cross-claim or
    counterclaim in a lawsuit) alleging that the Program itself (excluding
    combinations of the Program with other software or hardware) infringes
    such Recipient's patent(s), then such Recipient's rights granted under
    Section 2(b) shall terminate as of the date such litigation is filed.

    All Recipient's rights under this Agreement shall terminate if it fails
    to comply with any of the material terms or conditions of this Agreement
    and does not cure such failure in a reasonable period of time after
    becoming aware of such noncompliance. If all Recipient's rights under
    this Agreement terminate, Recipient agrees to cease use and distribution
    of the Program as soon as reasonably practicable. However, Recipient's
    obligations under this Agreement and any licenses granted by Recipient
    relating to the Program shall continue and survive.

    Everyone is permitted to copy and distribute copies of this Agreement,
    but in order to avoid inconsistency the Agreement is copyrighted and may
    only be modified in the following manner. The Agreement Steward reserves
    the right to publish new versions (including revisions) of this Agreement
    from time to time. No one other than the Agreement Steward has the right
    to modify this Agreement. IBM is the initial Agreement Steward. IBM may
    assign the responsibility to serve as the Agreement Steward to a suitable
    separate entity. Each new version of the Agreement will be given a
    distinguishing version number. The Program (including Contributions) may
    always be distributed subject to the version of the Agreement under which
    it was received. In addition, after a new version of the Agreement is
    published, Contributor may elect to distribute the Program (including
    its Contributions) under the new version. Except as expressly stated in
    Sections 2(a) and 2(b) above, Recipient receives no rights or licenses
    to the intellectual property of any Contributor under this Agreement,
    whether expressly, by implication, estoppel or otherwise. All rights in
    the Program not expressly granted under this Agreement are reserved.

    This Agreement is governed by the laws of the State of New York and the
    intellectual property laws of the United States of America. No party to
    this Agreement will bring a legal action under this Agreement more than
    one year after the cause of action arose. Each party waives its rights
    to a jury trial in any resulting litigation.
