## Supplementary material for "Paper-and-pencil questionnaires analysis: a new automated technique to reduce analysis time and errors": LightQuest_script_package: LightQuest_Manuscript.pdf

### Abstract

**Background and Objective:** Questionnaires are essential tools in many scientific fields, including health and medicine. However, the analysis of paper-and-pencil questionnaires is time consuming, source of errors and expensive, limiting its use in large cohort studies. Computer-based questionnaires might be a valuable alternative but they may introduce bias, especially for sensitive questions, and they require programming skills. The aim of this study is to develop a reliable and adaptable open-source technique (i.e. LightQuest) to automatically analyse various types of scanned paper-and-pencil questionnaires with closed questions, including those with inverted scale.

**Methods:** To evaluate the usefulness of LightQuest, the time needed for 7 experimenters for manually code 10 sets of 4 frequently used questionnaires and the number of errors (i.e. reliability) were compared with the time and errors their made using LightQuest.

**Results:** LightQuest was twice as fast as the manual analysis, even though the time to create the reference model was taken into account (933s vs. 1935s, $t(2)=8.81$, $p<0.001$). Without model creation, the reduced analysis time was more pronounced, with an average of $2.77 s.question^{-1}$ for the manual technique versus $0.55 s.question^{-1}$ for LightQuest ($t(2)=22.5$, $p<0.001$). Moreover, during correction of the 5180 questions performed by the 7 experimenters, LightQuest made a total of 2 errors versus 46 with the manual technique ($q(2)=4.53$, $p<0.05$).

**Conclusion:** LightQuest demonstrated clear superiority both in terms of time and reliability. The script of this first open-source technique, which does not require programming skills, is downloadable in supplemental data and may become an asset for all studies using questionnaires.


**Keywords:** Open-source technique; Psychometric; Computerization; Closed-ended survey

**Abbreviations:**

AF : Automatic with Feedback

AnF: Automatic with no Feedback

GIGO: Garbage In/Garbage Out

MCQ: multiple choice question

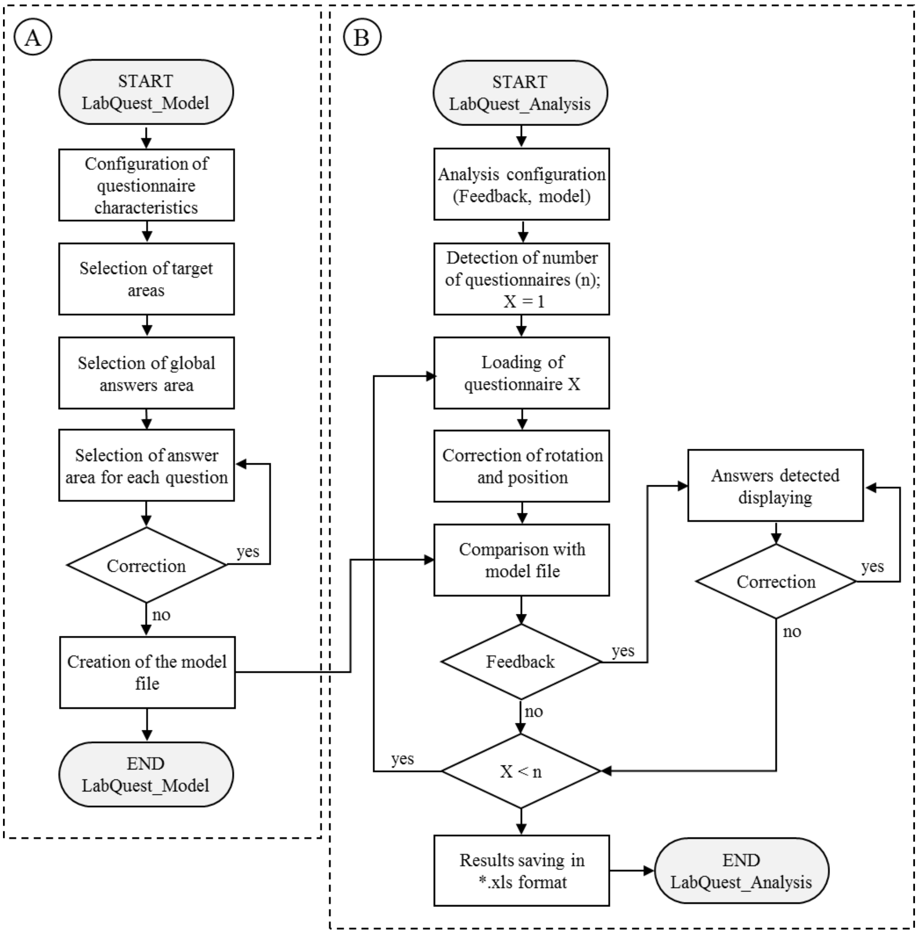


*<u>Figure 1:</u> Main steps of LightQuest processing for questionnaire model creation (A) and analysis by experimenters of the completed questionnaires (B).*

Second, the analysis of the questionnaires is performed by the LightQuest_Analysis.m file (Figure 1B). Once the script is run, a dialog box appears so that the appropriate model file can be selected to analyse the questionnaires, as well as the feedback level wanted during the analysis. If the feedback selected is “Yes”, each questionnaire is displayed with the detected answers, and the user can quickly verify and correct the analysis with an interactive interface in the case of multiple answers or no answer to a question (i.e: AF). Otherwise, selecting “No” means that the Excel matrix with the detected answers is directly generated by the script without steps of user verification and correction (i.e. AnF). For each participant of the study, 3 columns were implemented in the Excel file and saved in the Results folder. The first column corresponds to the detected answers, the second to the value attributed to the answer during model creation, and the third to the number of answers detected for each question. If inverted items are present in the questionnaire, the script automatically corrects the value attributed to the answer, in accordance with the configuration established during model creation. For ease of use, an explanation file describing the step by step use of the software is available in supplemental data (LightQuest.zip).

##### 2.5 Statistical analysis

The analysis time, detected answers, and number of errors were collected for each technique and questionnaire. Analysis time (in seconds) per questionnaire and per question were assessed with an omnibus 2-way ANOVA according to the questionnaire (PANAS, STAI, RSES, PANAS) and technique (manually, automatically with feedback: AF to directly validate and correct the detected answers, and automatically with no feedback: AnF). To perform the omnibus 2-way ANOVA, data were corrected with Box-Cox transformation [29] to fit with normality law (Shapiro-Wilk test) and homogeneity (Levene test). Holm-Sidak *post-hoc* tests were performed to determine the detected effects. Because results did not show significant differences between questionnaires (results not shown), statistical analysis of the total number of errors with each technique (Figure 3) was performed with a Kruskal-Wallis 1-way ANOVA on ranks (technique factor), followed by a Tukey *post-hoc* test.

### 3 Results

#### 3.1 Comparison of analysis times

Figure 2A presents the mean time taken by the 7 experimenters to analyse 10 copies of the PANAS, POMS, RSES and POMS questionnaires with the 3 techniques. Statistical analysis showed a main effect of both technique (F(2, 72) = 125.26, $p < 0.001$, partial $\eta 2 = 0.74$) and questionnaire (F(3, 83) = 4.66, $p = 0.005$, partial $\eta 2 = 0.041$), with no significant interaction between these 2 factors (F(6, 83) = $4.9e^{-4}$, $p = 0.812$, partial $\eta 2 = 8.7e^{-3}$).

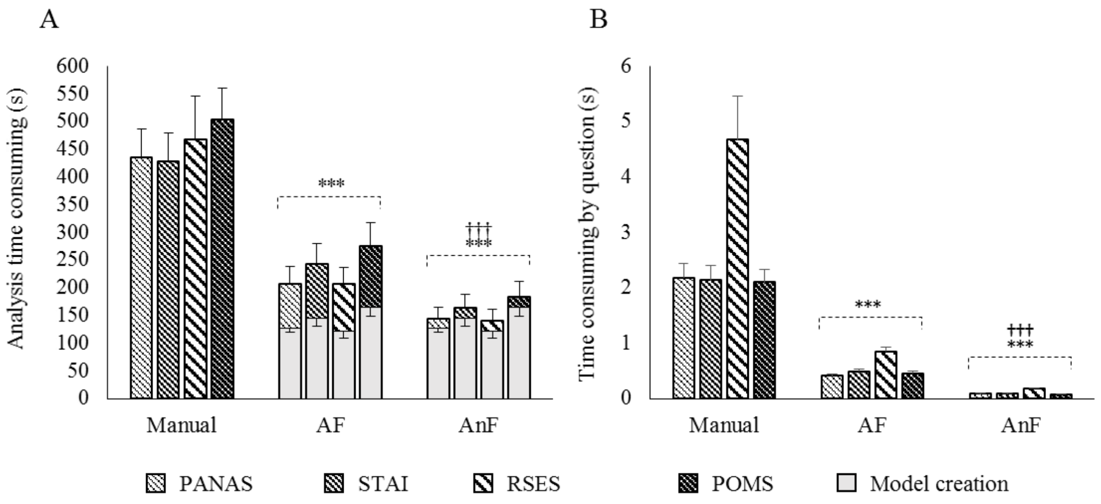


*<u>Figure 2:</u> Time needed for 10 questionnaires analysis (A) or for one question (B), according to the technique used. Mean±SEM; n = 7. AF: automatic + feedback; AnF: automatic with no feedback;* PANAS: *Positive and Negative Affect Schedule; STAI: State-Trait Anxiety Inventory; POMS: Profile of Mood States; RSES: Rosenberg Self-Esteem Scale. ***: diff. from manual analysis (p<0.001); †††: diff. from AF (p<0.001).*

Comparison of manual *versus* AF and AnF showed a significant decrease in analysis time between 49% and 66% (manual *vs*. AF: t(2) = 8.81, $p < 0.001$; manual *vs*. AnF: t(2) = 15.79, $p < 0.001$). Comparison of AF and AnF revealed a significant 32% decrease in analysis time by technique: the time in AF condition was significantly longer than the time in AnF condition ($t(2) = 6.98$, $p < 0.001$). *Post-hoc* analysis of the questionnaire factor showed a significant difference between POMS (24 questions) *versus* the RSES (10 questions) and PANAS (20 questions) questionnaires (respectively $t(3) = 3.35$, $p = 0.008$; and $t(3) = 3.06$, $p = 0.015$).

Analysis time in relation to the number of questions in each questionnaire (Figure 2B) also showed a main effect of technique ($F(2, 72) = 1083.42$, $p < 0.001$, partial $\eta 2 = 0.92$) and questionnaire ($F(3, 83) = 41.69$, $p < 0.001$, partial $\eta 2 = 0.053$), with no significant interaction between these 2 factors ($F(6, 83) = 0.57$, $p = 0.756$, partial $\eta 2 = 1.44e^{-3}$). Thus, the manual technique was 79% and 96% significantly slower than AF and AnF (respectively $t(2) = 22.50$, $p < 0.001$ and $t(2) = 46.54$, $p < 0.001$). AF and AnF were statistically different ($t(2) = 24.03$, $p < 0.001$), with the analysis time reduced over 80% with AnF compared with AF. Analysis of the questionnaire factor revealed that processing RSES was significantly longer than processing PANAS, POMS and STAI (respectively $t(3) = 9.42$, $p < 0.001$; $t(3) = 9.39$, $p < 0.001$; and $t(3) = 8.46$, $p < 0.001$).

##### 3.2 Reliability of the analysis techniques

To test the reliability of the analysis techniques, we compared the total number of errors made by each experimenter when they corrected the questionnaires manually and with AF and AnF (Figure 3).

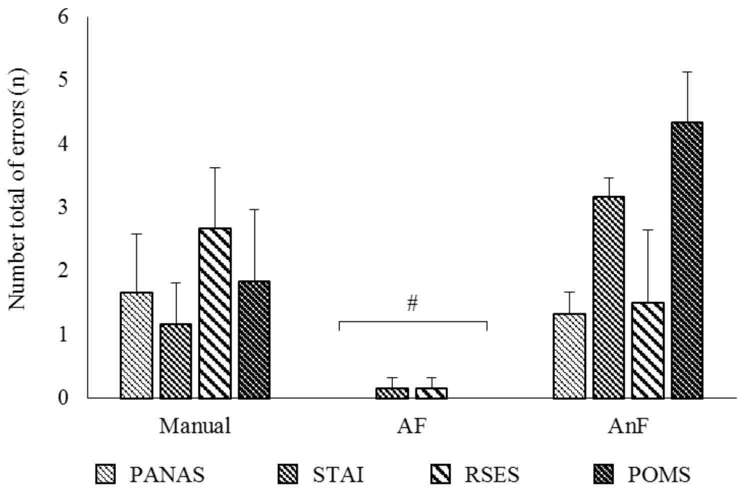


*Figure 3: Number of errors made by experimenters during analysis with the techniques. Mean±SEM; n = 7. AF: automatic + feedback; AnF: automatic with no feedback;* PANAS: *Positive and Negative Affect Schedule; STAI: State-Trait Anxiety Inventory; POMS: Profile of Mood States; RSES: Rosenberg Self-Esteem Scale. #: diff. from manual and AnF technique (p<0.05).*
