## Supplementary figures and images for "Paper-and-pencil questionnaires analysis: a new automated technique to reduce analysis time and errors"

### Model_POMS.jpg

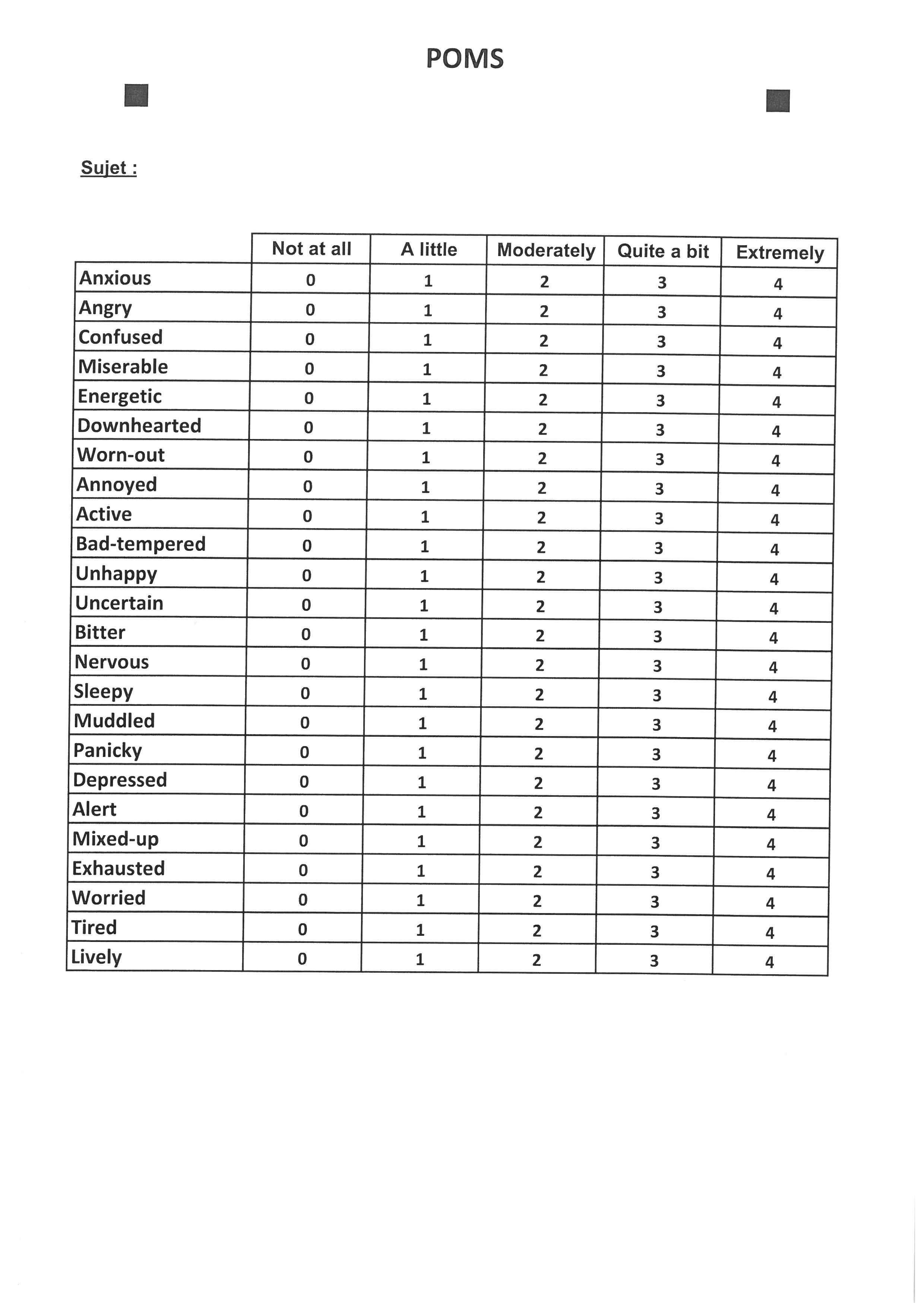

### S55C-6e17120510530_0001.jpg

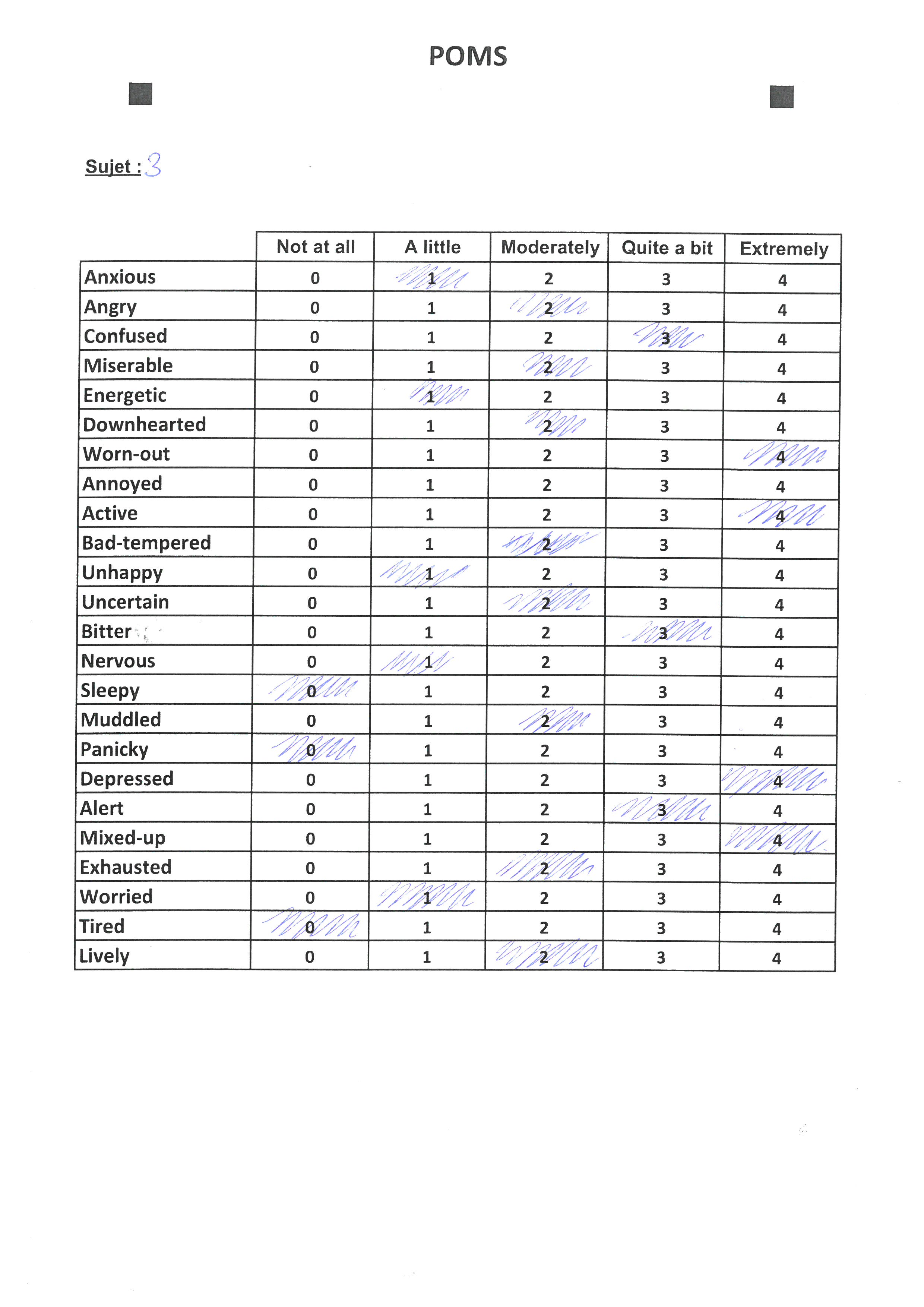

### S55C-6e17120510530_0002.jpg

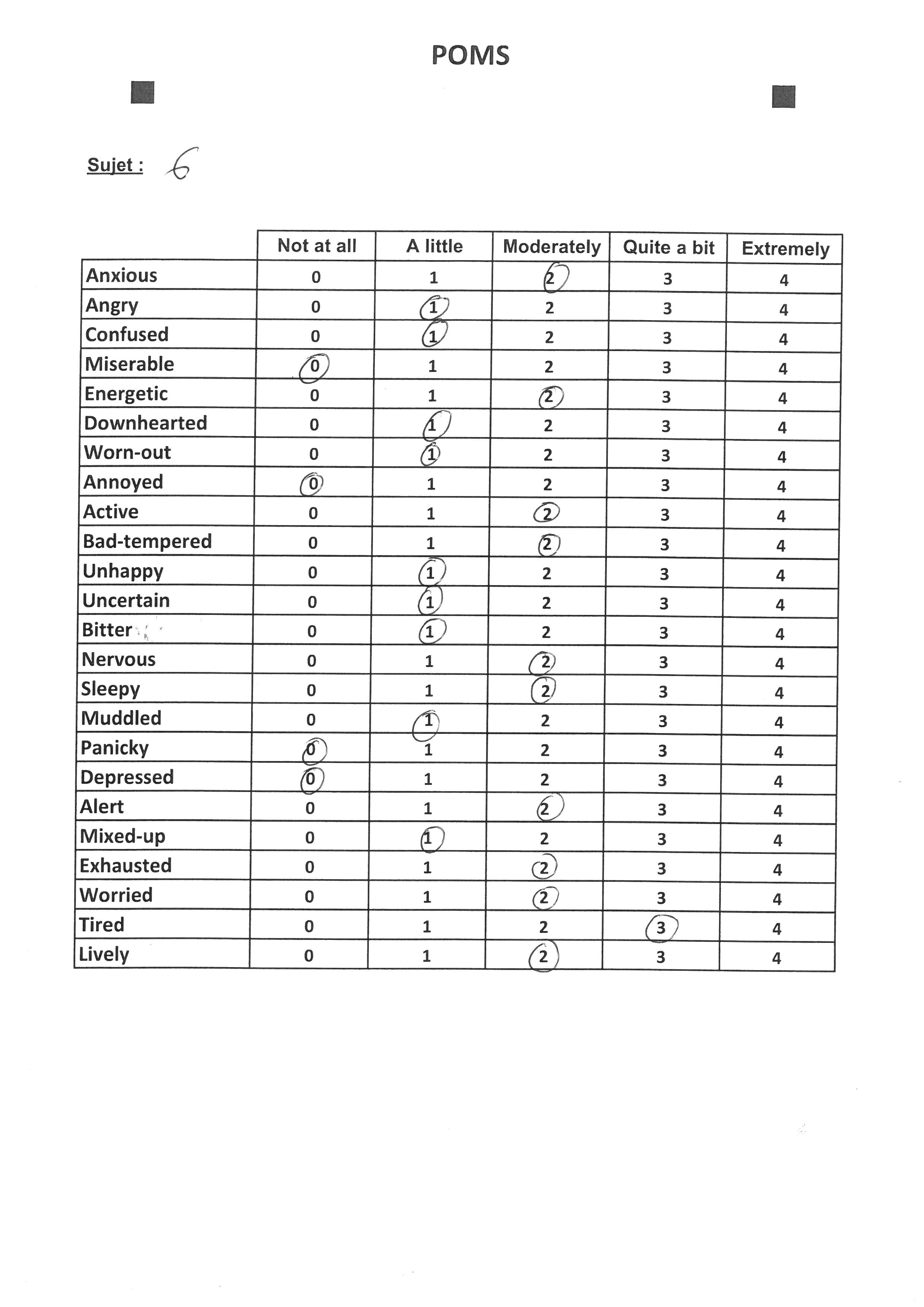

### S55C-6e17120510530_0003.jpg

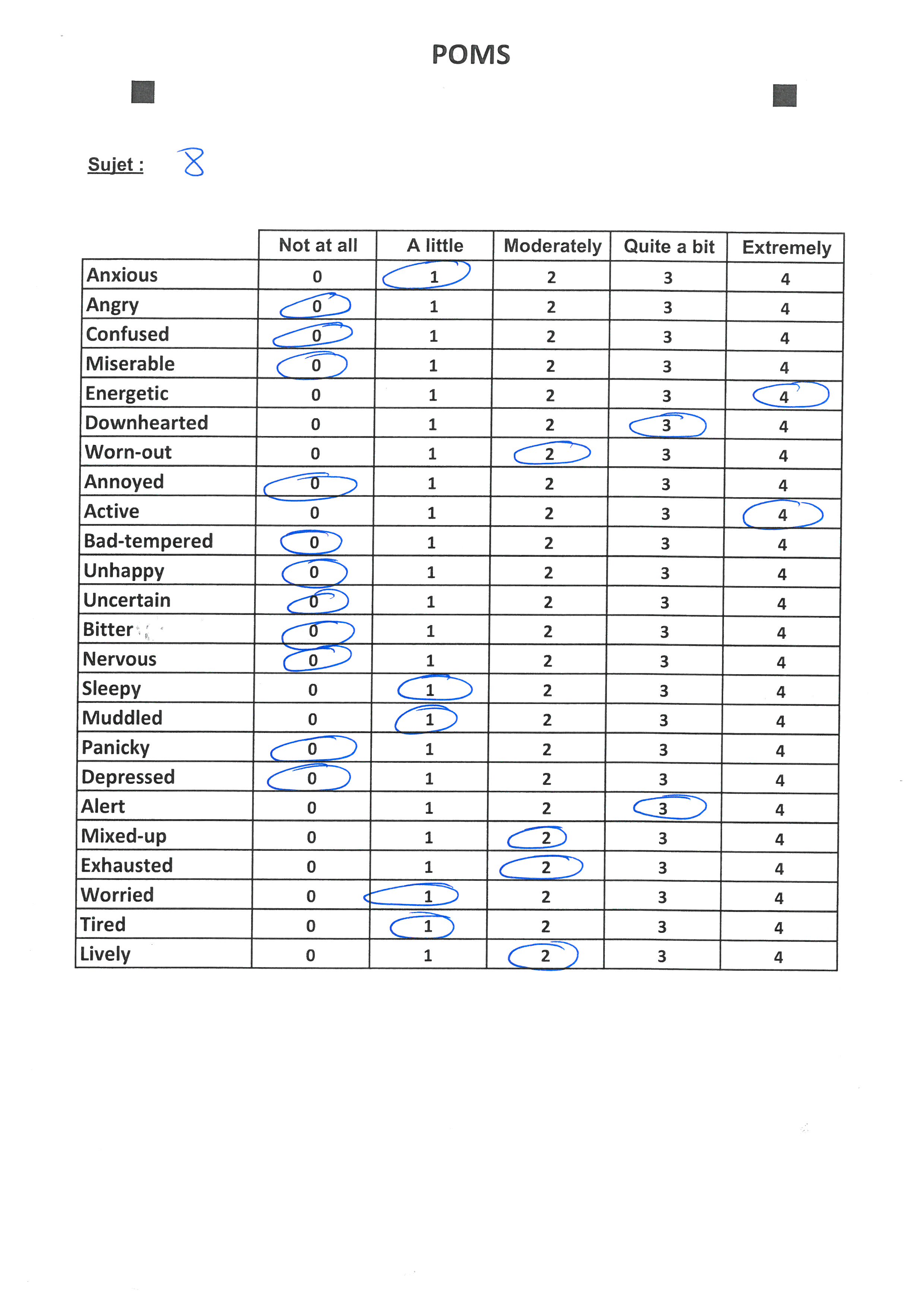

### S55C-6e17120510530_0004.jpg

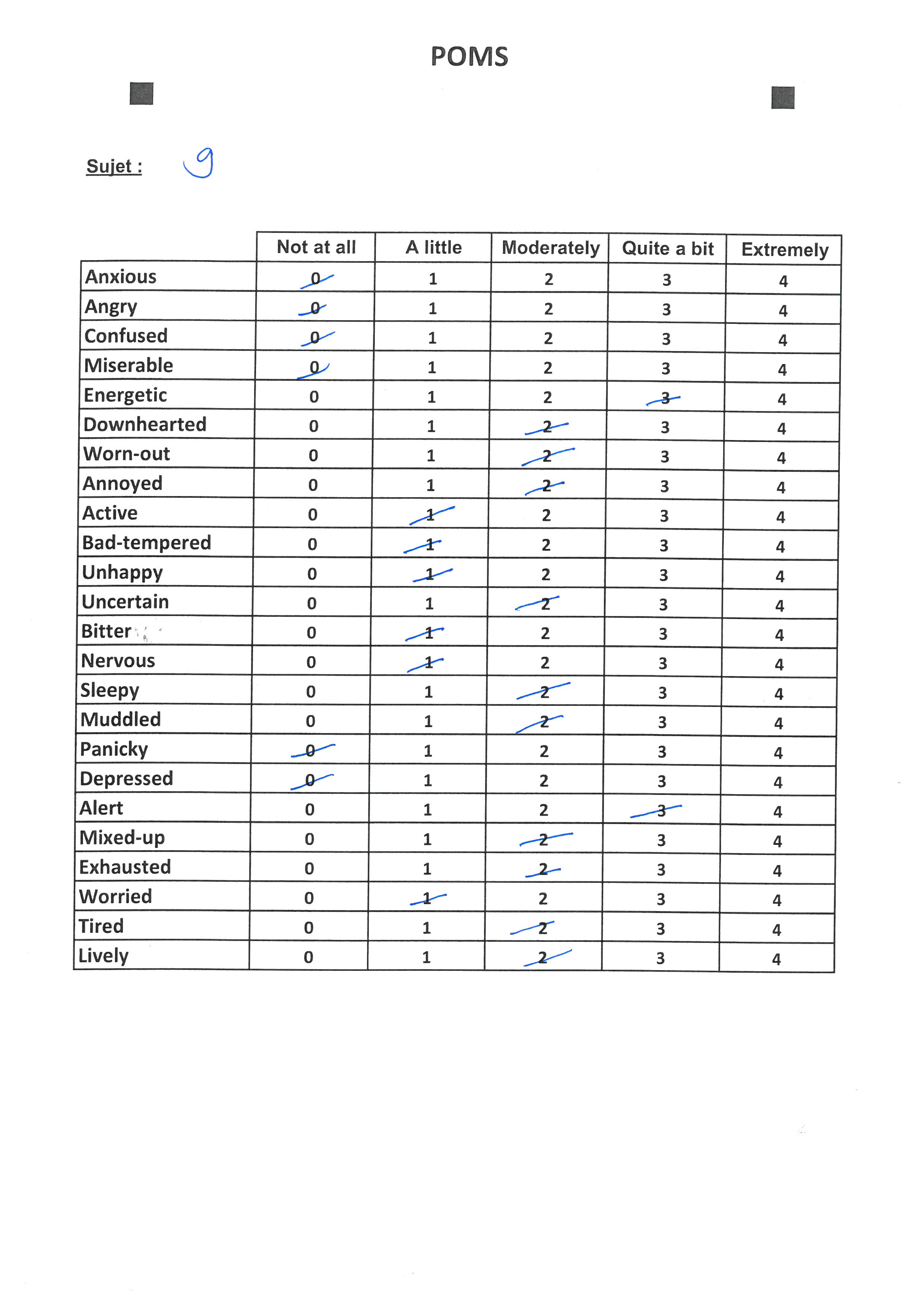

### S55C-6e17120510530_0005.jpg

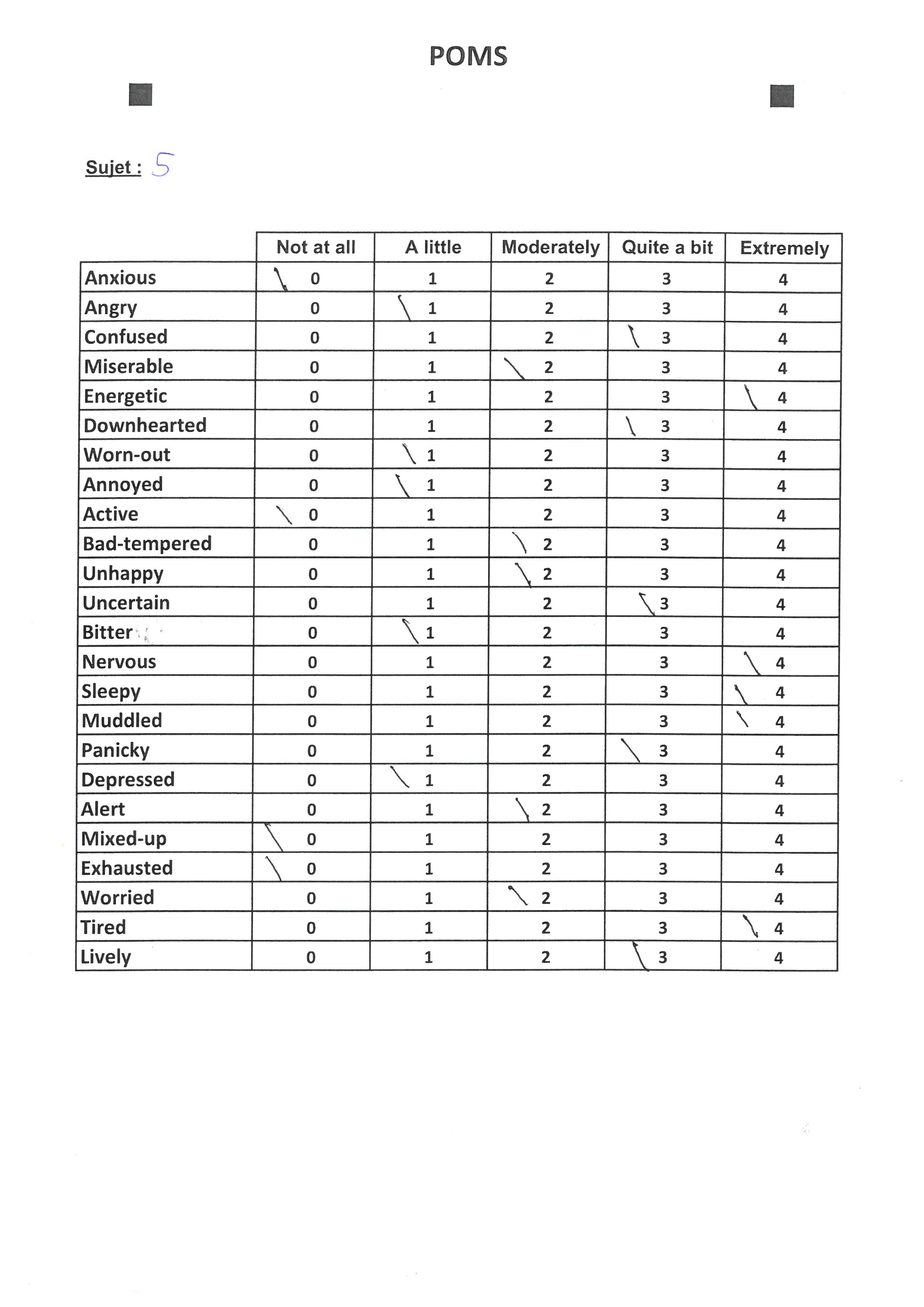
